# Mild uncoupling of mitochondria synergistically enhances senolytic specificity and sensitivity of BH3 mimetics

**DOI:** 10.1101/2023.08.23.554476

**Authors:** Edward Fielder, Abbas Ishaq, Evon Low, Joseph Laws, Aisha Calista, Jemma Castle, Thomas von Zglinicki, Satomi Miwa

## Abstract

Anti-senescence interventions are exceptionally effective in alleviating a wide range of age-associated diseases and disabilities. However, the sensitivity and specificity of current senolytic interventions are limited. Mitochondrial dysfunction is an integral part of the senescent phenotype and we demonstrate that specific loss of complex I-linked coupled respiration and the inability to maintain mitochondrial membrane potential upon respiratory stimulation are early and persistent features in a cell’s progression towards senescence.

We thus identify senescence-associated mitochondrial dysfunction as a targetable vulnerability of senescent cells and show that further decreasing mitochondrial membrane potential of senescent cells with a low concentration of a mitochondrial uncoupler synergistically enhances the *in vitro* senolytic efficacy of BH3 mimetic drugs, including Navitoclax, by up two orders of magnitude.

Moreover, in an *in vivo* mouse model of radiation-induced premature ageing, we show that a short-term intervention combining the mitochondrial uncoupler BAM15 with Navitoclax at a dose two orders of magnitude lower than typically used reduces frailty and improves cognitive function for at least 8 months after irradiation. Therefore our study shows that compromised mitochondrial functional capacity is a specific vulnerability of senescent cells which can be targeted by mild uncoupling *in vitro* and *in vivo*.

## Introduction

Together with apoptosis, cellular senescence is one of the major cellular stress responses with pleiotropic consequences during development and ageing. While cell senescence contributes to organ remodelling and wound healing, accumulation of senescent cells during ageing has been proven as a major cause of many, if not all, major age-associated diseases and disabilities, as specific ablation of senescent cells using senolytic drugs was able to postpone and in some cases even cure them ^1^. First-generation senolytics have been shown to be effective in alleviating many diseases and disabilities of ageing in mouse models and a growing number of clinical trials employing mainly the senolytics Dasatinib plus Quercetin (D+Q), or Fisetin, are underway ^2, 3^. However, one main limitation of current senolytics is a narrow therapeutic window, defined by the difference in EC50 between senescent and non-senescent cells, which is frequently less than one order of magnitude in concentration. This poses a significant risk for toxicity: for example, Navitoclax in concentrations typically used as senolytic (50mg/kg) can cause thrombocytopenia ^4, 5, 6, 7^, promote pulmonary hypertension ^8^ and impact bone mass in aged mice ^9^. The development of more specific senolytics is still an unmet clinical need.

Senescent cells differ from their non-senescent counterparts in many respects. In addition to a stable cell cycle arrest, they display major shifts in gene expression and the production and release of a host of bioactive molecules including cytokines, chemokines, pro-oxidative factors and other signalling molecules, termed the Senescence-Associated Secretory Phenotype (SASP). Senescent cells are also characterised by numerous changes in their mitochondria. Mitochondrial functional capacity is best described by the ability to homeostatically maintain the mitochondrial membrane potential (MMP). The MMP is generated by proton pumping by the electron transport chain using metabolic substrates, is consumed by processes such as proton leak, ATP synthesis and transport and is under constant homeostatic regulations. Senescent cells display low maximum respiratory capacity, increased proton leak and low MMP at basal state ^10, 11^ which together indicate compromised mitochondrial functional capacity. Further accompanying changes are increased mitochondrial mass, altered mitochondrial morphology, and increased generation of reactive oxygen species (ROS) ^11, 12, 13, 14, 15^. Together, these features have been termed Senescence-Associated Mitochondrial Dysfunction (SAMD) ^16, 17^. Mitochondrial functional alterations are observed during ageing in isolated mitochondria from various tissues including liver, skeletal and cardiac muscle and the brain. Literature shows that Complex I in the electron transport chain is the predominant site of the age-dependent dysfunction, with reduced coupling and increased ROS release with complex I linked substrates ^18, 19, 20, 21, 22^.

Interventions that reduce mitochondrial dysfunction in senescent cells have the potential to prevent or even reverse at least some components of the senescent phenotype. Complete ablation of mitochondria in senescent cells normalized cellular ROS levels, expression of cyclin-dependent kinase inhibitors (CDKIs) including p16 and p21, levels of senescence-associated β-galactosidase (Sen-β-Gal) and production of multiple SASP factors to levels similar to non-senescent cells ^23^. Similar suppression of the senescent phenotype was achieved by dietary restriction ^24^ which improves mitochondrial function (reviewed ^25^), and with dietary restriction mimetics such as rapamycin that reduce mitochondrial ROS ^23^.

Conversely, mitochondrial dysfunction in senescent cells may be proposed as a targetable weakness of senescent cells. In cells with compromised mitochondrial functional capacity, mitochondria are unable to maintain MMP upon increased ATP demand or other processes that consume MMP, and are more susceptible to prolonged mitochondrial permeability transition pore (mPTP) opening ^26^ and possibly, to (minority) mitochondrial outer membrane permeabilization (MOMP) ^27, 28^, both of which are linked to apoptotic cell death. Apoptosis triggered by MOMP is controlled by Bcl-2 family proteins and can be initiated in response to a plethora of intrinsic stress stimuli such as DNA damage, oxidative stress, and endoplasmic reticulum (ER) stress. Senescent cells are typically exposed to such stresses, and maintain high levels of anti-apoptotic Bcl-2 family proteins ^29^ to counter pro-apoptotic signalling and display resistance to apoptosis. BH3 mimetic drugs inhibit anti-apoptotic Bcl-2 family proteins, allowing the release of apoptogenic proteins including cytochrome c as well as Smac/DIABLO from the mitochondrial intermembrane space into the cytoplasm, which drives activation of the cascade of aspartate-specific cysteine proteases (caspases) that bring about apoptotic cell death ^30, 31, 32^. Suppressing the anti-apoptotic action of Bcl-2 proteins with BH-3 mimetics leaves the pro-apoptotic signalling in senescent cells unchecked, while non-senescent cells remain unaffected, thus selectively eliminating senescent cells and acting as senolytics ^33^.

Because senescent cells have a poor capacity to maintain MMP compared with non-senescent cells, we hypothesised that mild uncoupling of mitochondria can enhance sensitivity of BH3 mimetics such as Navitoclax to induce apoptosis specifically in senescent cells. Non-senescent cells on the other hand, would be able to tolerate mild uncoupling and thus would receive minimum effects under the same condition. We propose that the addition of low doses of mitochondrial uncoupler should enable the therapeutic efficacy of BH3 mimetics to be reached at substantially lower doses, hence broadening the therapeutic window and reducing the risk of side effects for senolytic interventions.

Here we first characterise the mitochondrial functional capacity of senescent cells and determine the concentration range of mitochondrial uncouplers that specifically targets senescent cell mitochondria. We then show that mild uncoupling enhances the senolytic activity of BH3 mimetics *in vitro* synergistically by up to two orders of magnitude without impacting non-senescent cells. Finally, we test the prediction that the MMP co-targeting approach enables to lower the effective doses of Navitoclax as senolytic in a mouse model of premature ageing *in vivo*.

## Results

### Senescence-associated mitochondrial dysfunction

To understand the development of senescence-associated functional changes in mitochondria, we performed a concerted time-course analysis of mitochondrial respiratory capacity, membrane potential and ROS levels following induction of stress-induced senescence (Fig. 1A-C). Human fibroblasts were treated with 20Gy irradiation, which is sufficient to induce senescence in 100% of the cells ^11, 12^. Mitochondrial oxygen consumption rates (OCR) using either complex I (Pyruvate and Malate) or II (Succinate) linked substrates were examined in permeabilised cells to determine mitochondrial coupled respiration. The respiratory control ratio (RCR) with complex I linked substrate was retained for at least 6 hours after irradiation but dropped by about 50% within the next 18 hours and then remained at low levels during further maturation of the senescent phenotype (Fig 1A), which was accompanied by a similar change in MMP (Fig. 1B). There was also a simultaneous increase in mitochondrial mass per cell as measured by MitoTracker® Green FM (MTG) fluorescence (Fig. 1B). On the other hand, complex II linked RCR remained largely unaffected (Fig. 1A). ROS levels in both mitochondria (MitoSOX™) and cytoplasm (DHE) only started to increase after complete reduction of complex I linked RCR and MMP at day 1 and continued to do so over at least one week (Fig. 1C). Thus in senescent cells, decreased coupled respiration at complex I is a key functional change and an early event preceding the increase of ROS.

**Fig. 1:**
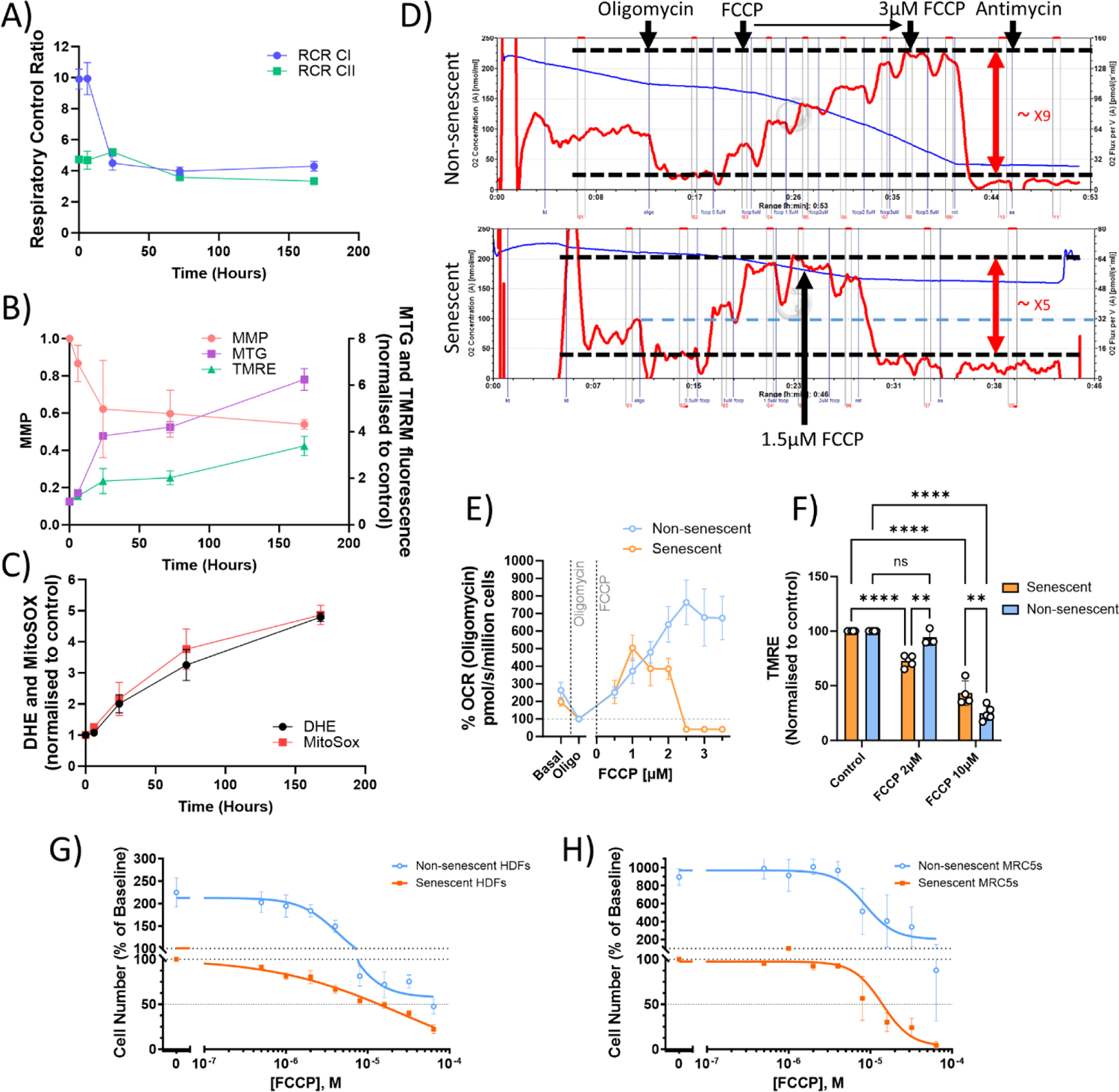
Mitochondrial functional changes during senescence development. A) – C) Senescence was induced in HDFs by 20Gy irradiation at time 0 and development of mitochondrial dysfunction was measured over the following 7 days. Time 0 indicates pre-irradiation. A) Respiratory Control Ratios (RCR) with complex I (blue) and complex II (pink) linked substrates. B) Mitochondrial mass (measured as MitoTracker® Green FM fluorescence intensity, green), TMRM fluorescence intensity (blue) and MMP (black). C) ROS production measured as DHE (red) and MitoSOX™ (yellow) fluorescence. D) Representative traces of intact cell oxygen consumption rates (OCR, red lines) measured in Oroboros O2k, with sequential additions of indicated compounds. FCCP was titrated by 0.5µM at each addition. Non-senescent cells can increase OCR about 9-fold compared to the resting state (in the presence of Oligomycin) whereas this is decreased to 5-fold in senescent cells. E) Summary of 6 (non-senescent) and 8 (senescent) independent Oroboros experiments. Values were obtained at steady-state after each addition of compounds, and presented as % of OCR at the resting state (in the presence of oligomycin). F) Changes in MMP in senescent (red) and non-senescent (blue) HDFs under the indicated concentrations of FCCP relative to control condition (no uncoupler). Data are mean ± SEM, N=3-8, *p≤0.05, **p≤0.005, ****p≤0.0005, ****p≤0.0001. G) and H) Change in number of senescent (red) and non-senescent (blue) HDFs (G) and MRC5 cells (H) after 3 days of treatment with the indicated FCCP concentrations. Data are mean ± SEM, N=3.

To further establish the link between low mitochondrial functional capacity and the ability to maintain MMP in senescent cells, and to explore the utility of mild uncoupling as a specific target for senescent cells, cellular OCR was measured with titrations of the mitochondrial uncoupler Carbonyl cyanide-p-trifluoromethoxyphenylhydrazone (FCCP) by 0.5µM up to 3.5µM in both senescent and non-senescent cells. As shown before by us and others ^10, 12^, ‘spare respiratory capacity’ (max OCR – basal OCR) was higher in non-senescent cells than in senescent cells; non-senescent cells tolerate uncoupling to more than 3µM resulting in about 8-fold higher maximum OCR compared with the resting state (in the presence of oligomycin), but respiration in the senescent cells collapses at an FCCP concentration beyond 2µM, achieving only about 5-fold higher maximum OCR compared to the resting state (Fig. 1D, E). 2µM FCCP was sufficient to reduce the MMP of senescent human fibroblasts, but not that of non-senescent fibroblasts. As expected, 10µM FCCP reduced MMP in both senescent and non-senescent cells (Fig. 1F). However, the uncoupler FCCP on its own in increasing concentrations did not lead to sufficiently differential reduction in cell viability between senescent and non-senescent cells. (Fig. 1G, H).

### Mitochondrial uncouplers synergistically increase the senolytic efficacy of BH3 mimetics

Having established the level of mild uncoupling that specifically targets MMP in senescent cells, we tested for a synergistic interaction between mild uncoupling and BH3 mimetics as senolytics.

We treated senescent and non-senescent human dermal fibroblasts (HDF) in co-culture with increasing concentrations of Navitoclax, which inhibits both Bcl-2 and Bcl-xL ^34^, with or without the mitochondrial uncoupler FCCP at 2µM and measured the change in cell number after 72 hours of treatment (Fig. 2A). By co-culturing HDFs constitutively expressing pSLIEW (containing EGFP) in stress-induced senescence (10 days past 20Gy irradiation, IR) with proliferating cells expressing mCherry and tracking changes in cell number by imaging, this model allowed us to take any interactions between senescent and non-senescent cells under the senolytics into account. In accordance with data from Fig. 1, 2µM FCCP had only a small effect on proliferation of non-senescent HDF during the 72 hours. However, numbers of surviving senescent HDFs were lower under all Navitoclax concentrations when combined with 2µM FCCP (Fig. 2A). Importantly, the combination with 2µM FCCP enhanced the EC50 of Navitoclax in senescent cells by about 2 orders of magnitude whereas it had a much smaller effect on non-senescent HDFs (Fig. 2B). Similar results were obtained for combinations of multiple mitochondrial uncouplers and BH3 mimetics: Navitoclax and CCCP (Fig. 2D), or the specific Bcl-xL inhibitor A1331852 and either FCCP (Fig. 2C) or CCCP (Fig. 2E).

**Fig. 2:**
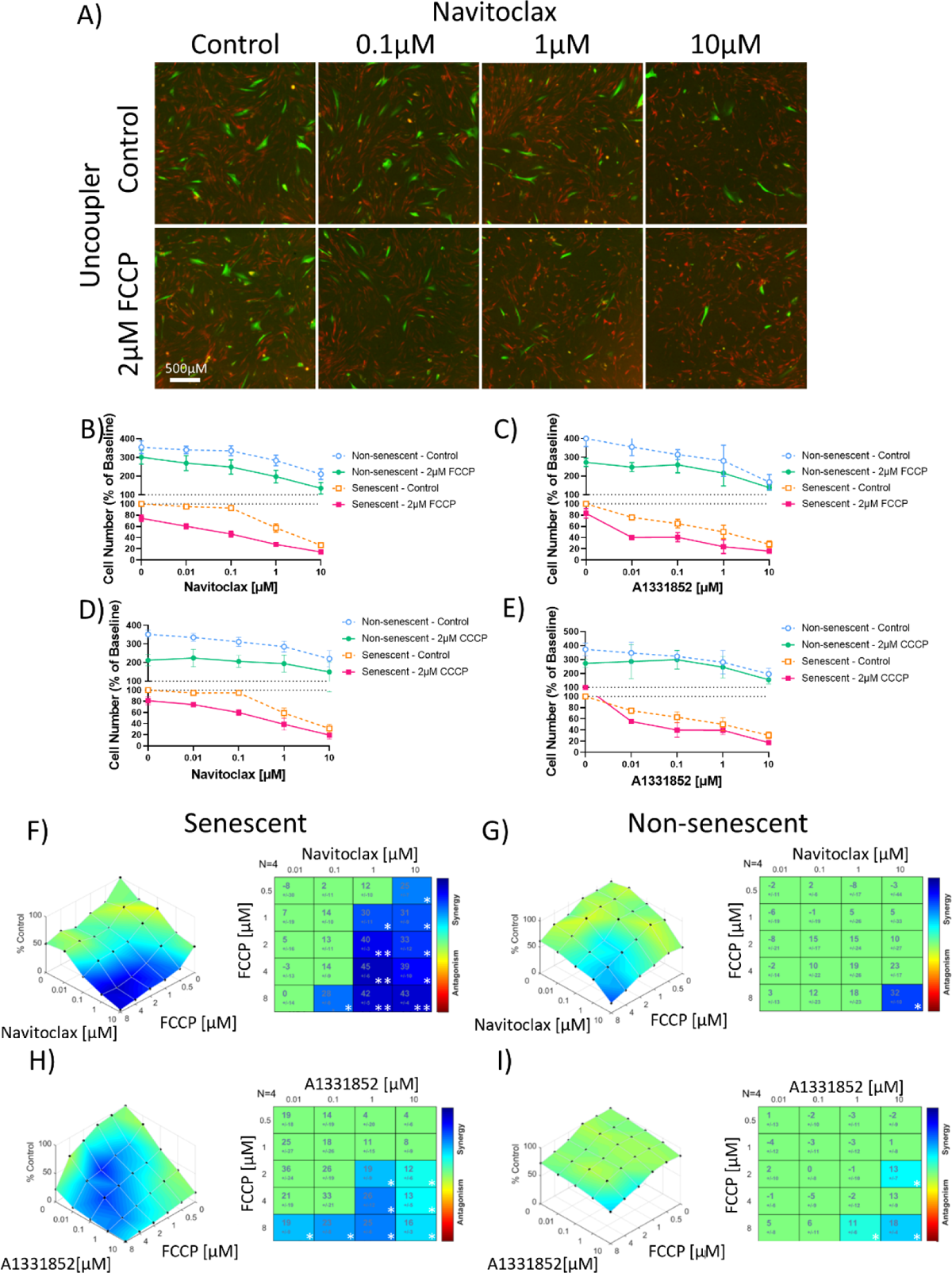
BH3 mimetics and uncouplers synergise in killing specifically senescent cells. A) Fluorescence micrographs of HDFs in X-Ray irradiation induced senescence (green) co-cultured with un-irradiated non-senescent HDFs (Red) for 3 days in the presence of increasing doses of Navitoclax with or without 2µM of FCCP. B) – E) Titration curves of senescent (red/yellow) and non-senescent (green/blue) HDFs with increasing concentrations of BH3 mimetic Navitoclax (B) and D)) or A1331852 (C) and E)) with or without 2 µM FCCP (B) and C)) or CCCP (D) and E)). Data are mean ± SEM, N=3-6 independent experiments. F) – I) Highest Single Agent analysis showing cell viability compared to control with combinations of BH3 mimetic and FCCP in senescent (left) and non-senescent cells (right). Synergy/antagonism is shown by colour and synergy score with change from expected value from either single compound for the Navitoclax/FCCP (F) and G) and A1331852/FCCP (H) and I) combinations on senescent (F) and H) and non-senescent (G) and I) HDFs. N=4, significance shown by * p≤0.05, ** p≤0.005.

To test for synergy between BH3-mimetics and mitochondrial uncouplers, we performed titration experiments using combinations of either Navitoclax or A1331852 and FCCP over a range of concentrations each. Highest single agent analysis using Combenefit showed strong synergism in senescent HDFs (Figs. 2F, H) but there was no significant interaction in non-senescent HDFs until the highest tested doses of both compounds (Figs. 2G, I).

To assess the mechanism of senolytic function of the combination of BH3-mimetics and mitochondrial uncouplers, cell viability, cytotoxicity and caspase3/7 activity were examined (Promega ApoTox-Glo Triplex Assay System) in senescent and non-senescent HDFs after 24 hours of treatment with Navitoclax alone (Suppl. Fig. S2.1A) or combined with 2µM FCCP (Suppl. Fig. S2.1B). There was very little induction of caspase activity in non-senescent cells, however, the induction of caspase activity by Navitoclax in senescent HDFs was potentiated by its combination with FCCP (Suppl. Fig. S2.1 A, B). Although a low concentration of Navitoclax (0.1µM) enhanced caspase activity in both non-senescent and senescent cells, a significant interaction between Navitoclax and FCCP was only found in senescent cells (not shown). Readouts for cell viability and cytotoxicity were independent of increasing Navitoclax concentrations with or without FCCP (Suppl. Fig. S2.1A, B), and addition of the pan-caspase inhibitor z-VAD-FMK for 24 hours reduced senescent cell loss under Navitoclax alone or with FCCP (Suppl. Fig. S2.1C). Thus these data together indicate that the combination of Navitoclax and FCCP induces apoptotic but not necrotic cell death. The combination treatment triggered a significant opening of the mPTP after 3 hours of treatment, which was rescued by cyclosporin-A, an inhibitor of mPTP opening (Suppl. Fig. S2.1D-E).

Navitoclax in combination with FCCP and CCCP also showed higher senolytic efficacy than Navitoclax alone in replicatively senescent HDFs (Suppl. Fig S2.2A-B), and in irradiation and replicatively induced MRC5 embryonic lung fibroblasts (Suppl. Fig S2.2C-D).

To test the specificity of the combination of BH3 mimetics and mitochondrial uncouplers as senolytics, we first tested an uncoupler that does not selectively target the MMP. 2,4-Dinitrophenol (DNP) is a weak uncoupler of both mitochondrial and other cellular membranes and prevents the uptake of inorganic phosphates into the mitochondria, leading to accumulation of potassium and phosphate ^35, 36, 37^. DNP concentrations up to 40µM had only mild effects on the senolytic activity of Navitoclax (Suppl. Fig S2.3A) or A1331852 (Suppl Fig S2.3B) in HDFs. To confirm that a reduction in MMP, rather than increasing cellular energetic output, enhances the senolytic function of BH3 mimetics, Monensin was used in combination with Navitoclax. Monensin is an ionophore which selectively transports sodium cations across lipid membranes and increases cellular energy expenditure via both glycolytic and oxidative phosphorylation. Monensin synergistically increased the killing effect of Navitoclax in both senescent and non-senescent cells equally, negating any specificity for senescent cells (Suppl. Fig. 2.3C, D).

We next combined mitochondrial uncouplers with senolytics that target other anti-apoptotic pathways that are upregulated in senescent cells. The cardiac glycoside Digoxin reduces pH by inhibiting the Na^+^/K^+^-ATPase pump, depolarizing the plasma membrane and causes a disbalanced electrochemical gradient within the cell, to which senescent cells appear to be more susceptible ^38^. In HDFs, however, the therapeutic window between senescent and non-senescent cells for Digoxin was small and was not improved by addition of either CCCP or FCCP (Suppl. Figs. 2.4A, B). D+Q inhibits multiple tyrosine kinases, PI3K and serpines and has been identified as senolytic in fat and endothelial cells but with limited senolytic activity in human fibroblasts ^39^. While uncouplers FCCP and CCCP reduced numbers of surviving senescent cells in some experiments, this was independent of an interaction with Dasatinib alone (Suppl. Figs. 2.4C, D) or D+Q in combination (Suppl. Figs. 2.4E, F). The naturally occurring flavonoid Fisetin was reported to have mild senolytic activity in fibroblasts ^40, 41^. We did not see significant senolytic activity of Fisetin in our assay, and this was not improved by combination with FCCP (Suppl. Fig. 2.4G). We conclude that only combinations of BH3 mimetics with mitochondrial uncouplers synergistically enhance senolytic sensitivity and specificity in fibroblasts.

It had been shown previously that chronic exposure to FCCP can have negative effects in human fibroblasts including induction of premature senescence ^42^. BAM15 is a recently discovered mitochondrial protonophore with a longer-lasting effect on respiratory kinetics and with improved safety profile that has been demonstrated *in vitro* and *in vivo* ^43, 44, 45^. Although chronic exposure to 2µM FCCP over 10 days reduced the growth rate of HDFs (Suppl. Fig. 3.1A) and induced senescence as shown by increased fractions of cells displaying karyomegaly (Suppl. Fig. S3.1B) and expressing Sen-β-Gal (Suppl. Fig S3.1C, D), 10µM BAM15 did not. When tested for its effect on cell viability on its own, the 10µM range of BAM15 could remove a fraction of senescent cells. (Suppl. Fig. S3.2A-F). Overall, BAM15 had slightly higher EC50 for fibroblasts than FCCP (Suppl. Fig. S3.2A-F, compare with Fig. 1), but with only a narrow difference between senescent and non-senescent fibroblasts (Suppl. Fig. S3.2A, D). Navitoclax combined with BAM15 killed stress-induced and replicatively senescent, but not non-senescent HDFs synergistically, from adult (Fig 3A-B, Suppl. Fig. S3.3A-B) and neonatal donors (Fig 3C-D), as well as MRC5 cells (Suppl. Fig. S3C-D). Replicatively senescent cells showed particular vulnerability to the combination (Suppl. Fig. S3.3B, D) as did chemotherapy-induced senescent MRC5 cells (Suppl. Fig. S3.3E).

**Fig. 3:**
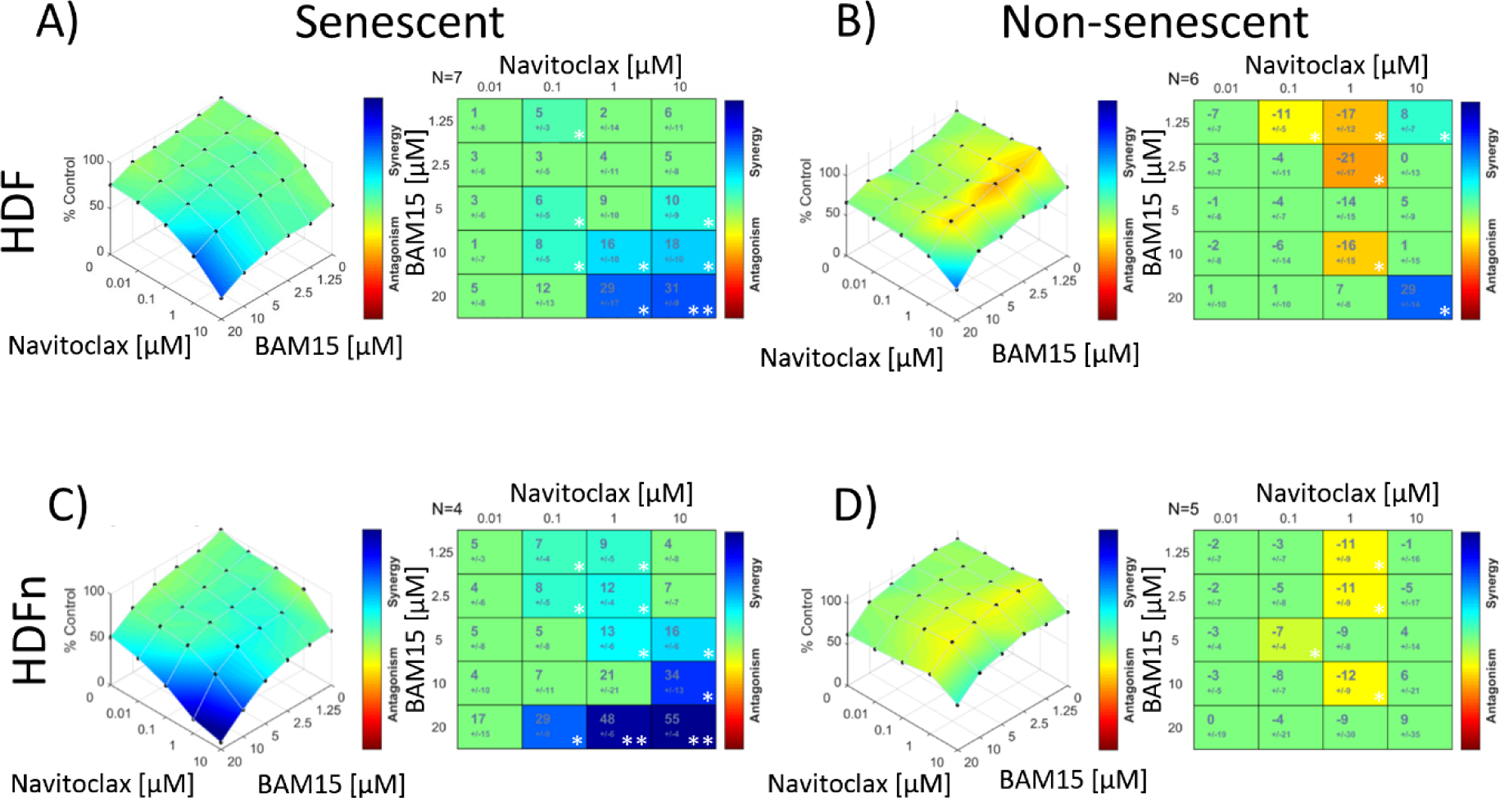
The combination of Navitoclax and BAM15 is synergistic for senolytic activity. Highest Single Agent analysis showing cell viability compared to control with combinations of BH3 mimetic and BAM15 in senescent (left) and non-senescent cells (right). Synergy/antagonism is shown by colour and, synergy score with change from expected value from either single compound for senescent and non-senescent adult HDFs (A, B) and neonatal HDFs (C, D) for BAM15 and Navitoclax, N=4-6. Significance shown by * p≤0.05, ** p≤0.005

10µM BAM15 enhanced Navitoclax-induced caspase activation in senescent but not non-senescent fibroblasts, and it did not induce necrosis (Suppl. Fig. S3.4A). Furthermore, 3 hours of combination treatment with Navitoclax and BAM15 induced more mPTP opening than Navitoclax alone, which was rescued by co-administration of cyclosporin-A (Suppl. Fig. S3.4B-C).

Cultured cells in nutrient rich- and high glucose-media can readily rely on glycolysis when mitochondrial ATP generation is compromised, and a typical cell culture environment can fail to recapitulate the *in vivo* environment where mitochondria generate almost all the energy in the form of ATP. Use of galactose as a replacement to glucose in culture media, which enhances mitochondria-driven ATP generation over glycolysis, often better reveals the *in vivo* scenario of drug toxicities around mitochondria ^46^. When tested in galactose media, the concentration of the compounds required for a synergistic effect in stress induced senescent HDFs was reduced (Suppl. Fig. S3.5).

Taken together, our data show that the mitochondrial uncoupler BAM15 also synergistically enhances the senolytic activity of Navitoclax in human fibroblasts.

It is widely hypothesized that therapy-induced senescence of tumour and niche cells can be a significant cause of insufficient tumour therapy response, and that this might be improved by adjuvant senolytic intervention past radio/chemotherapy ^47, 48, 49, 50, 51^. Tumour cells frequently show evidence of mitochondrial dysfunction even without senescence induction. We therefore tested the efficacy of the combination of mitochondrial uncoupler and Navitoclax on the human leukaemia cell lines 1301 and HL60, and the mouse glioblastoma line CT-2A. BAM15 alone was effective at arresting the growth of proliferating and therapy-treated leukaemia and glioblastoma cells (Suppl. Fig. S3.6A-D) and potentiated the effect of Navitoclax against both replicating and therapy-treated cancer cells (Suppl. Fig. S3.6A-C) with a synergistic interaction (Suppl. Fig. S3.6D). Senescent cancer cells can show resistance to Bcl-xL inhibition through upregulation of another anti-apoptotic Bcl-2 family member, Mcl-1 ^52, 53^. Using the Mcl-1 inhibitor (S63845) potentiated the effect of Navitoclax against therapy treated CT-2A cells, and worked in combination with BAM15 to further increase cell killing by another order of magnitude (Suppl. Fig. S3.7A). This effect could also be observed in stress-induced senescent MRC5 cells (Suppl. Fig. S3.7B). However, Mcl-1 inhibitors have dose limiting toxicities that could limit their use, and we found that at higher doses the combination of Navitoclax and S63845 was toxic to non-senescent fibroblasts (Suppl. Fig. S3.7C).

### In combination with the mitochondrial uncoupler BAM15, very low concentrations of Navitoclax are sufficient to rescue premature ageing in mice

Navitoclax is well established as an effective senolytic drug capable of alleviating multiple ageing- and/or stress-associated degenerative conditions if given in concentrations between 5 and 50 mg/kg BW to mice (Suppl. Fig. S4.1) ^54, 55, 56, 57, 58, 59, 60, 61, 62, 63^. In a model of irradiation-induced premature ageing ^64^ we established previously that male C57Bl/6J mice that received fractionated whole-body irradiation at an age of about 6 months developed frailty at about twice the normal ageing rate and showed decreased neuromuscular and cognitive function ^64^, all of which were rescued by a short course of 5mg/kg BW Navitoclax at one month after irradiation ^57^. Here, we employed the same model to test the efficacy of the combination of low Navitoclax with BAM15 *in vivo*. In addition to the previous dose of Navitoclax at 5mg/kg/day as in ^57^, we now treated the mice with either a 10-fold lower dose of Navitoclax at 0.5mg/kg/day, BAM15 at 2.5mg/kg/day, or with a combination of low Navitoclax at 0.5mg/kg/day and BAM15 at 2.5mg/kg/day (Figure 4 A). Treatment was for two rounds of 5 days, separated by 2 days. This design allowed us to address two questions in parallel, namely whether the combination was better than solo treatments alone, and how it compared to an intervention with the previously used higher Navitoclax doses.

**Fig. 4.**
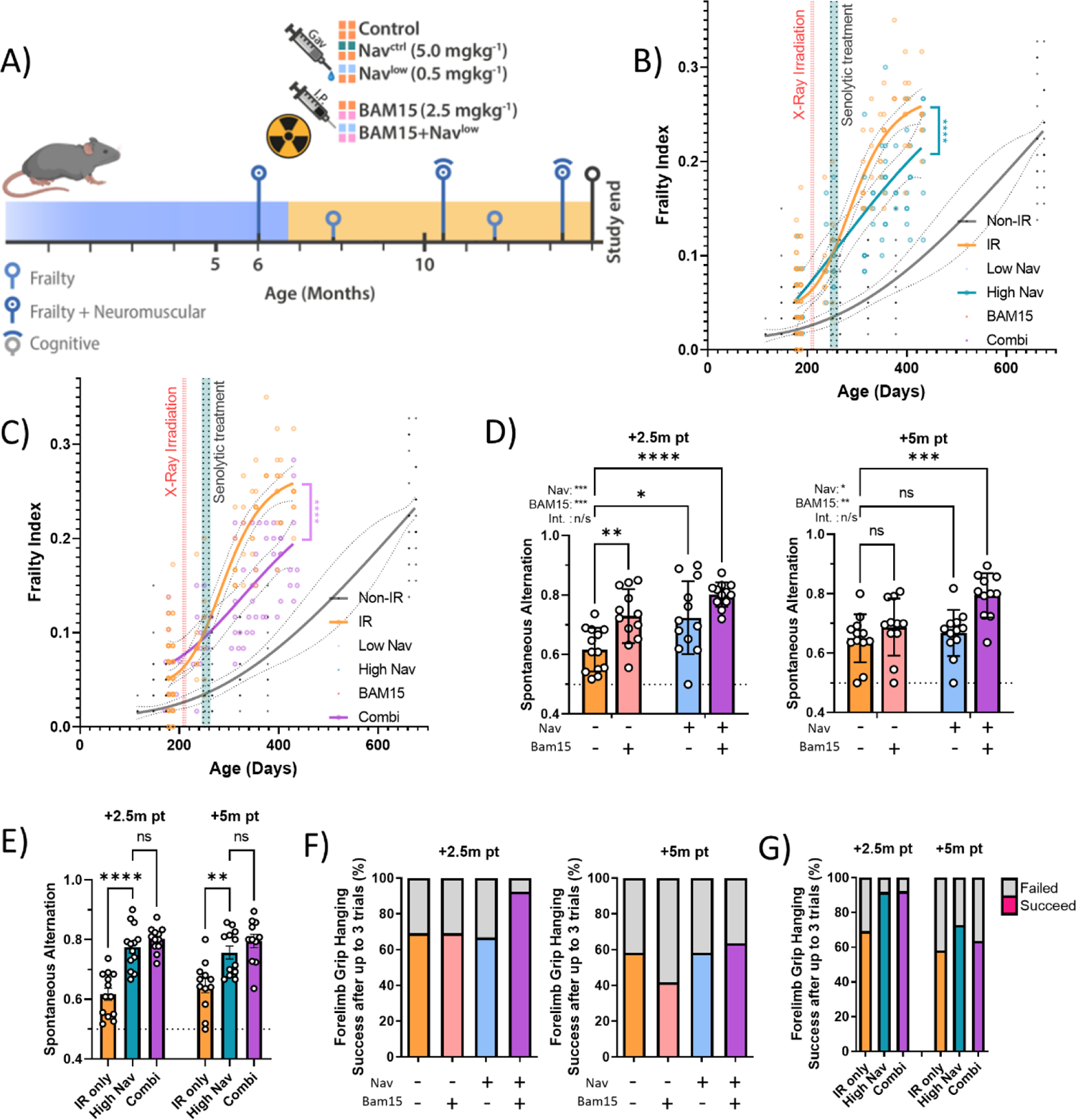
In combination with BAM15, a low dose of Navitoclax effectively rescued accelerated frailty and memory loss in irradiated mice. A) Experimental design. Mice were irradiated at 6 months of age and treated 1 month later with either vehicle, low or high dose of Navitoclax (0.5 and 5mg/kg respectively), BAM15 (2.5mg/kg) or a combination of low dose Navitoclax and BAM15. Mice were longitudinally phenotyped for frailty, cognition and neuromuscular performance over further 7 months. B) Frailty index (FI) vs mouse age for non-irradiated (Non-IR, black), irradiated (IR, orange), and irradiated plus treated with high Navitoclax (blue) mice. C) Frailty index (FI) vs mouse age for non-irradiated (Non-IR, black), irradiated (IR, orange), and irradiated plus treated with the combination of low Navitoclax plus BAM15 (pink) mice. B) and C) Irradiation and treatment times are indicated by vertical lines. Dots indicate FI for individual mice, regression lines and 95% confidence intervals are indicated by bold and dotted lines, respectively and variable slope (4 points) non-linear regression analysis is shown. Non-irradiated data is reproduced from Fielder *et al.* 2019 ^64^ for comparison. D) Short-term memory as assessed by spontaneous alternation in a Y maze in irradiated mice treated with either sham, BAM15, low Navitoclax or the combination of both at 2.5 months (left) and 5 months (right) past treatment. Comparisons by 2-way ANOVA with Šídák’s multiple comparisons post-hoc test against the untreated irradiated control, *≤0.05, ** p≤0.005, *** p≤0.0005, **** p<0.0001. Data are mean ± SEM, N≥11. E) Short-term memory as assessed by spontaneous alternation in a Y maze in irradiated mice treated with either sham, high Navitoclax or the combination of BAM15 and low Navitoclax at 2.5 months (left) and 5 months (right) past treatment. Data are mean ± SEM, N≥11. Comparisons by 1-way ANOVA with Šídák’s multiple comparisons post-hoc test against the untreated irradiated control, *≤0.05, ** p≤0.005, *** p≤0.0005, **** p<0.0001. F) Success rate in the Hanging Wire test for irradiated mice treated with either sham, BAM15, low Navitoclax or the combination of both at 2.5 (left) and 5 months (right) months past treatment. N≥11. G) Success rate in the Hanging Wire test for irradiated mice treated with either sham, high Navitoclax or the combination of BAM15 and low Navitoclax at 2.5 months (left) and 5 months (right) past treatment. N≥11.

None of the short-term treatments had a lasting effect on bodyweight (Suppl. Fig. S4.2A). Irradiated mice showed an early increase in frailty after irradiation and this was ameliorated by high Navitoclax (5mg/kg, Fig. 4B), as found before ^57, 64^. A low dose of Navitoclax or BAM15 alone resulted only in minor reductions of frailty progression (Suppl. Figs. S4.2B, C). However, the combination of BAM15 with a low Navitoclax dose was as effective in reducing frailty as a 10-fold higher dose of Navitoclax alone (Fig. 4C, compare to Fig. 4B).

Low Navitoclax, BAM15 and their combination improved cognitive function when assessed by measuring spontaneous alternation in a Y-maze at 2.5 months after treatment (Figure 4D) in 2-way ANOVA analysis. In a post-hoc analysis, only the combination treatment remained significantly different from controls at a later time-point, i.e. 5 months after treatment (Fig 4D). The higher dose of Navitoclax at 5mg/kg/day also significantly improved cognitive performance at both time points, to the same degree as the combination intervention (Fig. 4E).

Assessment of neuromuscular coordination by hanging wire test showed initial improvements by the combination treatment over low Navitoclax or BAM15 alone, although the effects were lost at the late time-point (Fig. 4F). An equal degree of improvement over irradiated controls were seen between the higher Navitoclax and the combination treatment, but only for the earlier time-point (Fig. 4G). Physical endurance, at either early or late time points after treatment, as measured by accelerating rotarod was not changed by any treatment (Suppl Fig S4.2D,F) and neither was liver damage as assessed by the activity of alanine transaminase (ALT) and aspartate aminotransferase (AST) in serum (suppl. Fig. 4.2E,G).

Given the persistent improvements in memory by the combination treatment (Fig. 4D-E), we examined markers of neuroinflammation in the hippocampus.

We measured the density of microglia (Fig. 5A, B) and their soma size (Fig. 5C, D) as a proxy of microglia activation in the CA2-3 region of the hippocampus, the dentate gyrus (Fig. 5E-H) and the periventricular zone (Fig. 5 I-L), and found reductions of both parameters after the combination treatment that were significant in most cases. 2-Way ANOVA indicated significant effects for BAM15 on microglia density (all examined regions) and for microglia soma size (Dentate gyrus and periventricular zone). Reductions in both parameters by the combination of a low dose of Navitoclax and BAM15 were not significantly different than those by the higher dose of Navitoclax alone (Figs. 5B, D, F, H, J, L). In accordance with a reduction in neuroinflammation markers, we also observed enhanced density of doublecortin-(DCX-) positive neurons in the subventricular zone as a proxy indicator of improved neurogenesis (Figs. 5M, N), which was significant in a post-hoc analysis for the combination treatment. Again, 2-way ANOVA analysis indicated a significant effect for BAM15 (Fig. 5M), and low Navitoclax together with BAM15 tended to be even more effective than the higher Navitoclax (Fig. 5N). While DCX positive cells were not common in the Dentate Gyrus, BAM15 showed a significant effect on Dentate Gyrus width, and post-hoc tests showed a significant change both alone and in combination with Navitoclax (Fig. 5O), which was not significantly different from the higher dose of Navitoclax (Fig. 5P). Together, our data show that a short intervention with a combination of a low dose of Navitoclax and BAM15 is sufficient to reduce late premature frailty, persistent neuroinflammation and cognitive impairment caused by sublethal irradiation as much as a high dose of Navitoclax.

**Fig. 5:**
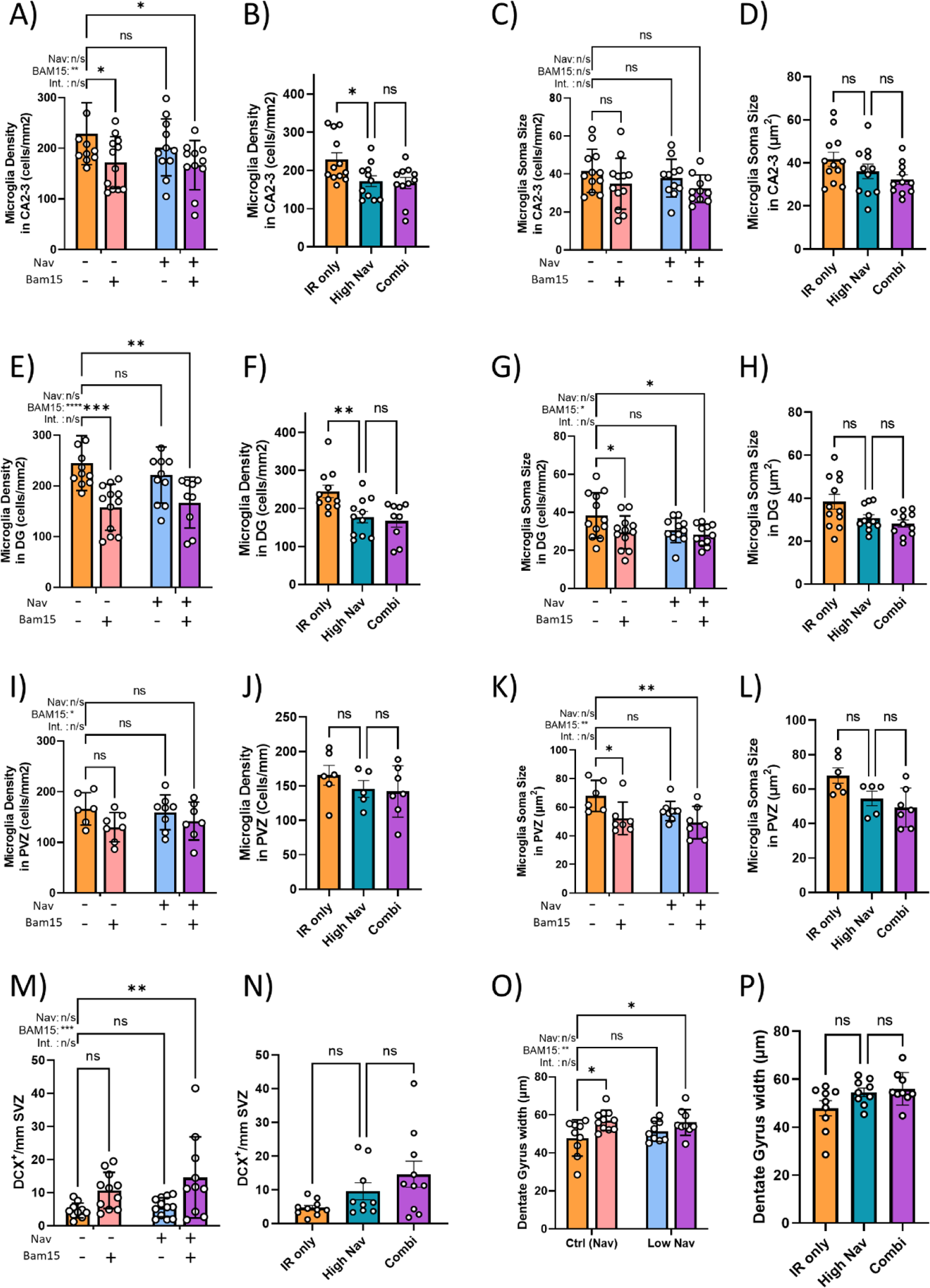
In combination with BAM15, a low dose of Navitoclax effectively reduced neuroinflammation at 7 months past treatment. Microglia density and soma size in the A-D) CA2-3 region of the hippocampus, E-H) the dentate gyrus and the I-L) subventricular zone in irradiated mice treated with either sham, BAM15, low Navitoclax or the combination. M-N) Numbers of DCX-positive cells along the subventricular zone. O-P) Thickness of the Dentate Gyrus. Data are mean ± SEM, sections from N=5-12 animals. 2-way ANOVA with Šídák’s multiple comparisons post-hoc for Control, BAM15, Low Nav and Combination. Comparisons by 1-way ANOVA with Šídák’s multiple comparisons post-hoc test against the untreated irradiated control for Control/High Nav/Combination, *≤0.05, ** p≤0.005.

## Discussion

Our kinetic study of mitochondrial functional changes during damage induced senescence *in vitro* showed decreased complex I dependent coupled respiration and decreased MMP as the earliest and persistent events preceding the increase in ROS production. Complex I dependent changes typically characterise mitochondrial functional changes in aged tissues, suggesting a mechanistic similarity between cell senescence and tissue ageing. Our study provides supportive evidence that mitochondrial dysfunction observed in senescent cells *in vitro*, as its limited ability to maintain MMP in response to mild uncoupling, is also functionally relevant *in vivo*: the mitochondrial uncoupler BAM15 was able to synergise with Navitoclax such that the same beneficial effects as 10 times higher dose were achieved in a mouse model of premature ageing.

The experiments on cells in culture were performed in an artificial condition, e.g. nutrient-rich media with high glucose. Under such conditions, senescent cells can compensate for their compromised mitochondrial functional capacity by shifting towards a more glycolytic energy metabolism. However, *in vivo* where oxidative metabolism is more dominant, it can be expected that the consequences of mitochondrial dysfunction in senescent cells could be even more serious. This could be partially mimicked by replacing glucose-with galactose-based media, which we found reduced the required concentrations for synergy of BAM15 and Navitoclax against senescent cells.

An important implication of the existence of populations of cells harbouring mitochondria with low ability to maintain MMP *in vivo* should be noted. Drug conjugates with a triphenylphosphonium (TPP^+^) lipophilic cation are frequently used in mitochondrial targeting approaches ^65, 66, 67^ because they enable rapid, several-hundred-fold accumulation of the conjugate into mitochondria in response to MMP ^65^. Importantly, such strategies typically aim to target ‘diseased’ cells, where mitochondria frequently exhibit compromised functional capacity and a correspondingly lower MMP. Examples include approaches to target cancer and senescent cells with cytotoxic drugs (Mito-Tamoxifen) or cancer cells with Mito-metformin ^68, 69, 70^. However, because of the low MMP in senescent or cancer cells, TPP^+^ conjugated drugs might preferentially concentrate in normal mitochondria in healthy cells *in vivo,* rather than in senescent or cancer cells, which can lead to unexpected outcomes. Conversely, TPP^+^ conjugated drugs may be more productive if mitochondria in non-senescent (or non-cancerous) cells are targeted, for example in order to protect them against adjuvant cytotoxic interventions. For example, mitochondria targeted antioxidants (MitoVit-E, MitoQ) have been shown to bring protective effects against oxidative stress induced damage, such as telomere shortening ^71, 72^.

There was a clear synergism between mitochondrial uncouplers and BH3 mimetics in senescent, but not non-senescent fibroblasts, with the combination having an up to 100-fold higher specific senolytic activity. This was achieved specifically by mitochondrial uncouplers and not by other drugs that cause generic upregulations of oxygen consumption rates and metabolic outputs. Furthermore, there was no senescence-specific synergistic activity in combinations of mitochondrial uncouplers with senolytics that do not directly target anti-apoptotic Bcl-2 family proteins. Our data suggested that the inability of senescent cells to maintain MMP under mild uncoupling significantly enhances their sensitivity to BH3 mimetics possibly through mPTP opening and/or MOMP. While the mechanistic and molecular links between MMP, mPTP opening and MOMP are still not fully understood ^22, 27, 28, 73^, it has been reported that mitochondrial uncouplers promoted Smac/DIABLO release from mitochondria to cytosol mediated by mPTP opening ^74^.

Our findings have important implications for better management of toxicity of senolytics *in vivo*. In combination with the uncoupler BAM15, a 0.5mg/kg/day dose of the BH3 mimetic Navitoclax was sufficient to rescue radiation-induced premature frailty and cognitive function in mice. Typical doses of Navitoclax used in mice are in the order of 50mg/kg/day (Suppl Fig. S4.1), indicating that a dose reduction by two orders of magnitude can be achieved *in vivo* by combination with an uncoupler. In clinical trials of patients with lymphoid tumours, at least 40% of patients reported thrombocytopenia at a dose of about 3-4mg/kg/day ^5, 6^. This is similar to what has been reported in rodents, and higher doses of Navitoclax have shown increased pulmonary hypertension ^8^ and bone loss in aged mice ^9^. Given that in rodents, human equivalent drug doses have been suggested to be around 12 times as high ^75^, a corresponding dose reduction, especially if only given for a short period (in our study, 2x 5 days was sufficient for an efficient and long lasting reduction), could potentially result in significantly less side effects. Among mitochondrial uncouplers, BAM15 has an excellent safely profile *in vivo*, appears to cross the blood brain barrier as it accumulated in brain amongst other organs ^45^, and has shown a potential in treating metabolic disorders and protecting organs from damage ^43, 44, 45, 76, 77^.

Our results may not only be relevant in the context of anti-ageing interventions, but also with respect to tumour chemotherapy, and for senolysis of both cancer and non-cancer cells in therapy-induced senescence. It is well established that many cancer cells show mitochondrial dysfunction with low MMPs and thus rely primarily on glycolysis, and drugs that lower MMP have been developed ^78^. We show that therapy-induced senescent cancer and non-cancer cells can also be (further) sensitised by mitochondrial uncouplers.

A limitation of our study is that cell type-specific senolytic activity was not comprehensively addressed. Generally, senolytics are active only in some cell types but not in others. This can be rectified to some extent by combining senolytic drugs as in the case of Dasatinib (senolytic for preadipocytes but not HUVECs) and Quercetin (senolytic for HUVECs but not preadipocytes) ^39^. We have established senolytic activity and specificity in multiple types of fibroblasts after different senescence inducers as well as in leukaemia and glioma cells *in vitro*. While the specificity (or not) for a wider spectrum of cell types still needs to be addressed, our *in vivo* results showing rescue of such complex phenotypes as frailty and short-term memory by the combination of low Navitoclax and BAM15 are encouraging. It is possible that reduction of senescence in fibroblasts across various organs could have benefitted tissue microenvironments, as fibroblast are a predominant cell type in stroma.

We have previously found that both D+Q and Navitoclax resulted in similar beneficial outcomes for multiple ageing phenotypes in parallel experiments *in vivo*, despite these drugs having different cell type specificities ^57^. In fact, recent results indicate that local, cell-type specific senolysis can be less effective than systemic senolysis even in the targeted organ ^79^. Beneficial effects of senolysis on brain function are thought to be largely due to systemic reduction of senescent cells and their SASP ^80^. Accordingly, senolytic drugs which are optimized for systemic rather than cell type-specific targeting could bring a wide range of benefits *in vivo*, and the BH3 mimetic and mitochondrial uncoupler combinations might be strong candidate leads.

## Methods

### Cell culture procedure

All cells were grown in a controlled environment of 5% CO_2_ and ambient oxygen at 37°C in Dulbecco’s Modified Eagle’s Medium (DMEM, Sigma Aldrich, D5671), supplemented with 10% heat-inactivated Foetal Bovine Serum (FBS, Sigma Aldrich, F9665), 100U/ml penicillin and 100mg/ml streptomycin (Sigma Aldrich, P4333), and 2mM L-Glutamine (Sigma Aldrich, G7513). 1301 and HL-60 cells were cultured in RPMI 1640 with 2mM L-Glutamine and 10% FBS.

Galactose media is prepared with 5.5mM D-(+)-Galactose (Sigma Aldrich, G0750) replacing glucose in DMEM (no glucose, no glutamine, no phenol red) (ThermoFisher, A1443001), phenol red (Sigma Aldrich, P3532) and 10mM HEPES (Sigma Aldrich, H0887), 10% heat-inactivated Foetal Bovine Serum (FBS, Sigma Aldrich, F9665), 100 U/ml penicillin and 100mg/ml streptomycin (Sigma Aldrich, P4333), and 2mM L-Glutamine (Sigma Aldrich, G7513).

Cell culture involved Human Dermal Fibroblasts (HDF) derived from the foreskin of an 8-year-old male donor, and neonatal HDFs sourced from neonatal foreskin (C0045C, Invitrogen). Other cell lines used included MRC5s (human foetal lung fibroblasts (ECACC 05011802)), CT-2A (mouse glioma cells (SCC194, Merck)), 1301 (Human T-cell Leukaemia cells (ECACC 01051619)) and HL-60 (Acute promyelocytic leukaemia (ECACC 98070106)).

Senescence was induced in fibroblasts through two mechanisms: stress-induced senescence, exposure to X-Ray irradiation (20Gy at 225kV) 10 days prior to the experiment, and replicative senescence, by culturing until the replication rate dropped below 0.2 population doublings per week. 1301 and HL-60 cells were exposed to X-Ray irradiation (4Gy and 6Gy, respectively at 225kV) 10 days prior to the experiment. CT-2A cells required 5 days of 25µM Temozolomide (BioTechne Ltd, #2706), 2 days rest, followed by 5 days of X-Ray irradiation (2Gy at 225kV).

### Mitochondrial function and cellular ROS

Mitochondrial respiratory capacity was examined in permeabilised cells in a Seahorse XF24 Analyzer, using Plasma Membrane Permeabilizer (PMP, Agilent) to permeabilise the cells (1nM). The cells were plated on XF24 cell culture microplates and irradiated as above, and the experiments were performed at indicated time points after the irradiation using Pyruvate (10mM) and Malate (1mM) or Succinate (4mM) in the presence of rotenone (0.5µM) as respiratory substrates for complex I and II respectively. After the measurement of basal oxygen consumption rate (OCR), the following compounds were added sequentially: 4mM ADP (state 3), 1µM oligomycin (state 4), 4µM Carbonyl cyanide-p-trifluoromethoxyphenylhydrazone (FCCP, uncoupled rate) and 2.5µM antimycin A (complete inhibition of the electron transport chain). The respiratory control ratio (RCR) was calculated as state 3 divided by state 4 OCRs.

OCRs from intact cells were measured using Oroboros O2k. The cells grown in flasks were gently trypsinised, pelleted, resuspended in the culture media, and placed in the chamber for the experiments immediately. After the basal OCR is established, the following compounds were added sequentially: 1µM oligomycin, FCCP in stepwise titrations (0.5µM each to 3.5µM), 0.5µM rotenone and 2.5µM antimycin A.

Cellular and mitochondrial ROS levels were measured using 5µM Dihydroethidium (DHE, D1168, ThermoFisher Scientific) and 0.5µM MitoSOX™ (M36008,ThermoFisher Scientific) respectively, by flow cytometry. Mitochondrial mass was measured using 20nM MitoTracker® Green FM (M7514, ThermoFisher Scientific), and mitochondrial membrane potentials using 25nM Tetramethylrhodamine (TMRM, T668, ThermoFisher Scientific) also by flow cytometry. The cells were grown on 6-well plates and irradiated as above. On indicated time points, the media was replaced with fresh media containing above probes (or vehicle control) and incubated at 37°C for 30 min. Then the cells were gently trypsinised, resuspended with the respective media with the probes and subjected to flow cytometry measurement immediately.

### pSLIEW and mCherry HDF generation

Early passage cells were transduced with pSLIEW or mCherry encoding virus (whole cell localisation) to generate HDF-pSLIEW (Green, eGFP) and HDF-mCherry (Red, mCherry). Plasmid DNA was extracted from E. Coli culture under ampicillin selection using the Invitrogen PureLink HiPure Plasmid Maxiprep Kit (K210006), according to manufacturer’s instructions. DNA concentration was determined using a nanodrop spectrophotometer (ND-1000), and plasmid DNA was stored at 1μg/μl in 1X Tris-EDTA (TE) buffer at −20°C.

Restriction digests were performed to confirm the isolated plasmids using BAMHI, SalI and XbaI in NEBuffer 3.1 #B7203S (New England Biolabs), PstI and HindIII in NEBuffer Blue #B7002S (New England Biolabs) and EcoRI in Cut Smart Green #B7204S (New England Biolabs), with 5% enzyme concentration, with 5% plasmid DNA. The digests were carried out at 37°C using a thermal cycler (Veriti, ThermoFisher Scientific) for 3 hours, and confirmed with gel electrophoresis on a 0.8% agarose gel containing 0.005% cyber green peak green DNA binder (Peolab, 37-5099).

Hek 293T packaging cells were cultured and transfected with 12μg of equimolar plasmid DNA using the Lipofectamine 3000 reagent kit (Invitrogen, L3000001). 6% Lipofectamine 3000 in serum-free DMEM media was mixed 1:1 with diluted plasmid DNA (serum free media with 2% plasmid DNA and 4% P3000 reagent). The generated virus from Hek 293T cells was supplemented with 6μg/ml polybrene and added to the target cells for 12 hours at 37°C.

### Senolysis assay

HDF-pSLIEW were plated into the interior 60 wells of black, clear bottomed 96 well plates (Greiner Bio-one #655090) at a density of 1.5k cells per well. The 36 edge-wells were filled with media. After 24 hours, the wells were supplemented with 50µl of fresh media, and the plate was irradiated with 20Gy of X-Ray. The media was replaced after irradiation and then changed every 3 days for 9 days post-irradiation.

On day 9 post-irradiation, proliferating HDFs expressing mCherry were co-cultured with the irradiated senescent HDF-pSLIEW at approximately 3k cells per well. Following a 24-hour rest period, the media was changed from DMEM to Fluorobrite imaging medium (Sigma Aldrich, F9665) with 5% FBS, 100U/ml penicillin, 100mg/ml streptomycin, and 2mM L-Glutamine for baseline imaging.

Images were captured in pre-set locations in each well using a Leica DMi-8 with an automated stage. Media was then switched to normal media containing the compound of interest, with controls receiving the same amount of solvent (DMSO or Ethanol). After a set period (72 hours unless otherwise specified), the media was changed back to imaging media, and images were taken again.

Image files were manually analysed to provide cell counts for senescent (green) and non-senescent (red) cells. Data were expressed as the percentage change between baseline and day 3 imaging, and senescent cells normalised to their control.

### Cell viability

Cells were plated into the interior 60 wells of clear 96-well plates (Corning, 3596) and allowed to settle for 24 hours. The exterior 36 wells were filled with media. To induce senescence, cells were irradiated with 20Gy (225kV) and kept for 10 days, with media changes immediately after irradiation, and then every 3 days. To account for proliferation, a ‘baseline measurement’ for proliferating cells was taken by fixing a secondary plate immediately prior to administering the drugs in media to the other plates. The plate was washed with PBS, fixed with 2% PFA for 5 minutes, washed twice more and kept at 4°C. On the measurement day (72 hours later) the plate was taken from the fridge, allowed to warm to room temperature, and the PBS replaced with warmed media. The plate was then processed with the remaining plates. Plates were washed with PBS, and incubated in 0.2% Crystal violet/1% Ethanol solution for 10 minutes. Plates were then washed twice by immersion in a basin of tap water and dried. Once dry, 1% SDS was added and plates put on a plate-shaker. Absorbance was read at 590nm. Background was subtracted using the empty exterior wells. Senescent cells were expressed as a percentage of control, while the baseline measurement was used to calculate the percentage change in cell number of the proliferating cells.

Cell viability of cells in suspension (HL-60 and 1301 leukaemia cells lines) was assessed using trypan blue exclusion (Sigma Aldrich, #93595) and manually counted using a haemocytometer.

### ApoTox-Glo triplex assay

The ApoTox-Glo triplex assay (Promega, G6320) were performed in 96 well plates (Greiner Bio-one #655090) according to manufacturer’s protocol. Results were red using a fluorescent plate reader (FLUOstar Omega, BMG), with no-cell and untreated cell controls.

### Mitochondrial permeability transition pore assay

Mitochondrial permeability transition pore opening was analysed using the Calcein/Cobalt technique^81^. Cells were exposed to different test drugs in normal media for 3 hours, media removed and cells counterstained with Hoechst in normal media (8µM) for 7 minutes, then media with 10mM CoCl_2_ was added 1:1 to wells for 7 minutes. Media was removed, and Calcein-AM (1µM) was added with 5mM CoCl_2_ for 15 minutes to quench fluorescence outside of the mitochondria. Media was removed and replaced with CoCl_2_ (3µM) supplemented Fluorobrite imaging media (Sigma Aldrich, F9665; with 100U/ml penicillin, 100mg/ml streptomycin, and 2mM L-Glutamine) medium for imaging using Leica DMi-8. Average intensity was recorded per cell with an average of 400 cells per condition, per repeat. Intensity following 10µM Ionomycin was subtracted, and results normalised to control.

### Senescence-associated beta-galactosidase (Sen-β-Gal)

Cells were fixed for 5 min with 2% PFA in PBS-Mg before incubation with the staining solution as described ^82^ (PBS-Mg containing 1mg/ml X-gal, 5mM potassium ferrocyanide, and 5 mM potassium ferricyanide, pH 6) overnight at 37°C. Cells were co-stained with DAPI and imaged for brightfield and fluorescence (350_Ex_, 460_Em_) using a Leica DMi-8 Brightfield. Cells were quantified as positive and negative based on the presence of blue staining.

### Drug synergy analysis

Drugs and their combinations were prepared in a 96 well plate in normal media, with duplicate wells. The media on 96 well plates containing senescent or proliferating cells was then exchanged for drug containing media, and kept for 72 hours. Following this crystal violet staining was performed as above, with background signal subtracted from blank exterior wells. Duplicate wells were averaged and then values being normalised to control wells (with the same solvent concentration as drug containing wells). Synergistic/antagonistic drug interaction was analysed using highest single agent analysis with Combenefit ^83^. Synergy score is expressed as percentage of response at each combination of doses that is beyond that expected from the individual compounds.

### Animals

Male C57Bl/6J mice were bought past weaning from Charles River and were maintained in groups of 3 littermates in IVC cages, otherwise as described ^84^. Mice were fed standard pelleted food (CRM-P formulation rodent diet, SDS Diets). Cage enrichment included extra bedding and plastic domes for nesting, two cardboard tunnels per cage, rings and chain for climbing, and chewing blocks. Mice were kept on a staggered light-cycle, switching to dark cycle at 2:30pm, with daily and weekly weighing performed under red-light conditions after the time. Mice were tunnel handled unless otherwise required experimentally. Mice were identified by tail marking until microchipped during group allocation with IMI-500 Read Only Transponder (Plexx). The work was approved by the UK Home Office (PBDAFDFB0) and complied with the guiding principles for the care and use of laboratory animals. Mice were irradiated at 6-7 months of age, as previously described ^64^. Mice received 1% Baytril solution in drinking water for 2 days before, and 14 days after, to the start and end of irradiation, respectively. Intervention for weight loss post-irradiation was performed as in ^57, 64^.

### Treatment of mice with drugs

Mice were orally gavaged with either empty vehicle, 0.5mg/kg BW Navitoclax or 5mg/kg BW Navitoclax for 10 days total (5 days followed by 2 days recovery, then 5 days again). Compounds were prepared for oral gavage in 10% Polyethylene Glycol (PEG) 400, vehicle control mice were gavaged with 10% PEG400 only. Mice were simultaneously given intraperitoneal injection of either empty vehicle or 2.5mg/kg BW BAM15 in 40% PEG400. PEG400 (8074851000) was purchased from Merck, BAM15 (Cat. No. 5737) was purchased from Biotechne, and Navitoclax (A3007-APE-100mg) was purchased from Stratech. The mice were phenotyped one month post-irradiation as below. Mice were then ranked on frailty score (low to high), average time on rotarod (high to low) and body weight (high to low). An average of each rank was then taken, to get a combined ranked score. Mice were then assigned to each group according to the combined ranked scores to reduce variability of the pre-dosing, baseline phenotypic measures between the groups.

### Mouse phenotyping

Frailty was assessed using a 30-parameter index ^85^, with modifications as described ^64^. Assessment was performed during the light-cycle by two people who were blinded to the treatment. Scores were given independently by both assessors and then averaged. Rotarod, hanging wire and spontaneous alternation Y-Maze were performed as described ^64^ during the dark-cycle under red-light conditions. During the Y-maze, luminosity was 1.7lux ±0.1lux at the top of each arm and 0.1lux ± 0.1 at the bottom of the maze. Y-maze duration was 8 minutes. For the hanging wire, previously used ^64^ thick cotton bedding was substituted with 10cm of memory foam; this provided a softer landing and eased retrieval of the mouse and cleaning.

### Liver damage assay

Liver damage was assessed using the Alanine Transaminase Activity Assay Kit (Abcam, ab105134) and Aspartate Aminotransferase Activity Kit (Abcam, ab 105135) according to manufacturer’s instructions in a 96 well plate using a plate reader (FLUOstar Omega, BMG).

### Immunohistochemistry and Immunofluorescence

Paraformaldehyde (PFA)-fixed paraffin embedded tissue samples were cut from coronally embedded blocks and stained with primary/secondary antibodies as described in Table 1. Immunohistochemistry/fluorescence, microscopy and analysis were performed as otherwise described ^57, 86^. Microscopy for immunofluorescence and immunohistochemistry was performed using a DMi8 fluorescence microscope (Leica, Germany). Microglia density and soma size were measured as described ^57, 86^ to give iba1+ cells per mm^2^, and soma size (µm^2^). DCX positive cells were counted, and divided by the perimeter of the lateral ventricle to give DCX^+^ cells per mm. Dentate Gyrus width was measured every 120µm along the blades, an average of these measurements was then taken for each mouse. Images were analysed using the Fiji distribution of ImageJ2.

**Table 1.**
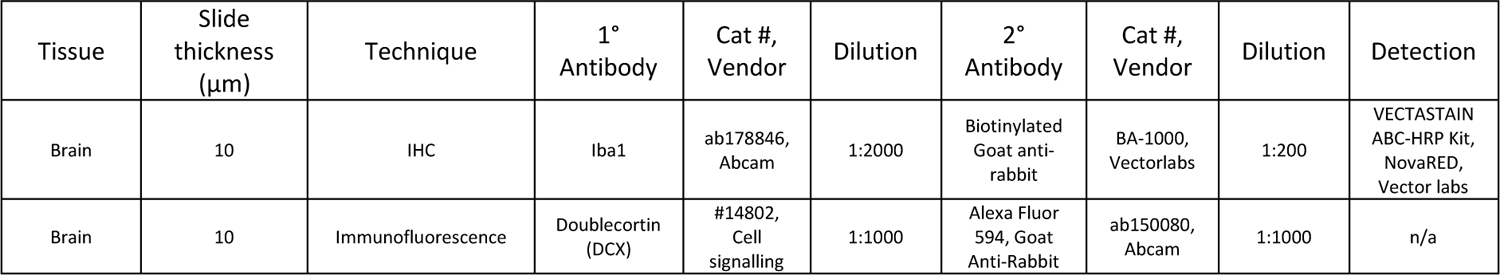
Conditions for IHC and IF.

### Navitoclax literature search

To survey the current dosing regimens using Navitoclax in the context of ageing and as an anti-senescence therapy, a literature for search on PubMed was performed as ((Navitoclax) OR (ABT-263)) AND (mouse). Papers ascertaining to senescence, senolytics and ageing in mice were selected from this list for inclusion (last updated 2023-04-04). Route of administration (Oral gavage, intraperitoneal, intramuscular, intravenous, intradermal and topical administration routes were identified), schedule of dosing (including total days) and doses used each time were manually extracted from each paper and tabled. Where this data was not clear from the paper, the authors were approached. If these data could not be obtained the papers were excluded from the results. As we were providing Navitoclax by oral gavage, papers using this route of administration were selected and total duration of administration and dose per dosing tabled as a .csv. A kernel density estimation plot these was created using Seaborn 0.11.1.

### Statistics

Data was collated in Microsoft Excel, with all statistics and graphing being performed in Graphpad Prism 9. One-way and two-way analysis of variance, and nonlinear regression analysis tests were performed using GraphPad Prism 9. Drug synergy was performed as described above.

The *in vivo* study was designed to evaluate if a lower (than previously administered by us) dose of Navitoclax (0.5mg/kg) would have an effect if combined with the mitochondrial uncoupler BAM15 (2.5mg/kg). Low Navitoclax and BAM15, alone and in combination, were compared to an untreated control group (vehicle only gavage and I.P.). This was set up as a two-way ANOVA and analysed in Graphpad Prism 9.

In order to assess if low Navitoclax and BAM15 in combination would be comparable to our previously used higher dose of Navitoclax (5mg/kg), the groups control, Nav (5mg/kg) and Nav (0.5mg/kg)/BAM15 (2.5mg/kg) were compared separately with a one-way ANOVA. To avoid repeating comparisons with the two-way ANOVA, the Nav/BAM15 group was not compared to the control in post-hoc for this test.

## Acknowledgements

We thank Christopher Huggins and the entire team at Comparative Biology Centre for their invaluable assistance and expertise during our experiments. Special thanks are also due to Naomi Buadee and Robyn Iredale for their exceptional technical support and unwavering commitment.

## References

1. Chaib S, Tchkonia T, Kirkland JL. Cellular senescence and senolytics: the path to the clinic. Nature Medicine 28, 1556–1568 (2022).

2. Sun Y, Li Q, Kirkland JL. Targeting senescent cells for a healthier longevity: the roadmap for an era of global aging. Life Medicine 1, 103–119 (2022).

3. Kirkland JL, Tchkonia T. Senolytic drugs: from discovery to translation. J Intern Med 288, 518–536 (2020).

4. Suvarna V, Singh V, Murahari M. Current overview on the clinical update of Bcl-2 anti-apoptotic inhibitors for cancer therapy. European Journal of Pharmacology 862, 172655 (2019).

5. Wilson WH, et al. Navitoclax, a targeted high-affinity inhibitor of BCL-2, in lymphoid malignancies: a phase 1 dose-escalation study of safety, pharmacokinetics, pharmacodynamics, and antitumour activity. The Lancet Oncology 11, 1149–1159 (2010).

6. de Vos S, et al. Safety and efficacy of navitoclax, a BCL-2 and BCL-XL inhibitor, in patients with relapsed or refractory lymphoid malignancies: results from a phase 2a study. Leukemia & Lymphoma 62, 810–818 (2021).

7. Schoenwaelder SM, et al. Bcl-xL-inhibitory BH3 mimetics can induce a transient thrombocytopathy that undermines the hemostatic function of platelets. Blood 118, 1663–1674 (2011).

8. Born E, et al. Eliminating Senescent Cells Can Promote Pulmonary Hypertension Development and Progression. Circulation 147, 650–666 (2023).

9. Sharma AK, et al. The Senolytic Drug Navitoclax (ABT-263) Causes Trabecular Bone Loss and Impaired Osteoprogenitor Function in Aged Mice. Front Cell Dev Biol 8, 354 (2020).

10. Hutter E, Renner K, Pfister G, Stöckl P, Jansen-Dürr P, Gnaiger E. Senescence-associated changes in respiration and oxidative phosphorylation in primary human fibroblasts. Biochem J 380, 919–928 (2004).

11. Passos JF, et al. Mitochondrial Dysfunction Accounts for the Stochastic Heterogeneity in Telomere-Dependent Senescence. PLOS Biology 5, e110 (2007).

12. Passos JF, et al. Feedback between p21 and reactive oxygen production is necessary for cell senescence. Mol Syst Biol 6, 347 (2010).

13. Moiseeva O, Bourdeau V, Roux A, Deschênes-Simard X, Ferbeyre G. Mitochondrial dysfunction contributes to oncogene-induced senescence. Mol Cell Biol 29, 4495–4507 (2009).

14. Mai S, Klinkenberg M, Auburger G, Bereiter-Hahn J, Jendrach M. Decreased expression of Drp1 and Fis1 mediates mitochondrial elongation in senescent cells and enhances resistance to oxidative stress through PINK1. Journal of Cell Science 123, 917–926 (2010).

15. Dalle Pezze P, et al. Dynamic Modelling of Pathways to Cellular Senescence Reveals Strategies for Targeted Interventions. PLOS Computational Biology 10, e1003728 (2014).

16. Korolchuk VI, Miwa S, Carroll B, von Zglinicki T. Mitochondria in Cell Senescence: Is Mitophagy the Weakest Link? EBioMedicine 21, 7–13 (2017).

17. Chapman J, Fielder E, Passos JF. Mitochondrial dysfunction and cell senescence: deciphering a complex relationship. FEBS Letters 593, 1566–1579 (2019).

18. Mansouri A, et al. Alterations in mitochondrial function, hydrogen peroxide release and oxidative damage in mouse hind-limb skeletal muscle during aging. Mechanisms of Ageing and Development 127, 298–306 (2006).

19. Cocco T, Pacelli C, Sgobbo P, Villani G. Control of OXPHOS efficiency by complex I in brain mitochondria. Neurobiology of Aging 30, 622–629 (2009).

20. Petrosillo G, Matera M, Moro N, Ruggiero FM, Paradies G. Mitochondrial complex I dysfunction in rat heart with aging: critical role of reactive oxygen species and cardiolipin. Free Radical Biology and Medicine 46, 88–94 (2009).

21. Miwa S, et al. Low abundance of the matrix arm of complex I in mitochondria predicts longevity in mice. Nature Communications 5, 3837 (2014).

22. Miwa S, Kashyap S, Chini E, von Zglinicki T. Mitochondrial dysfunction in cell senescence and aging. The Journal of Clinical Investigation 132, e158447 (2022).

23. Correia-Melo C, et al. Mitochondria are required for pro-ageing features of the senescent phenotype. The EMBO Journal 35, 724–742 (2016).

24. Wang C, et al. Adult-onset, short-term dietary restriction reduces cell senescence in mice. Aging (Albany NY) 2, 555–566 (2010).

25. Miwa S, Johnson TE. Stress in Aging. In: Molecular and Cellular Biology of Ageing (eds Vijg J, Campisi J, Lithfow G). GSA Press, 216–249 (2016).

26. Bernardi P. Modulation of the mitochondrial cyclosporin A-sensitive permeability transition pore by the proton electrochemical gradient. Evidence that the pore can be opened by membrane depolarization. J Biol Chem 267, 8834–8839 (1992).

27. Renault Thibaud T, et al. Mitochondrial Shape Governs BAX-Induced Membrane Permeabilization and Apoptosis. Molecular Cell 57, 69–82 (2015).

28. Ichim G, et al. Limited mitochondrial permeabilization causes DNA damage and genomic instability in the absence of cell death. Mol Cell 57, 860–872 (2015).

29. Ryu SJ, Oh YS, Park SC. Failure of stress-induced downregulation of Bcl-2 contributes to apoptosis resistance in senescent human diploid fibroblasts. Cell Death & Differentiation 14, 1020–1028 (2007).

30. Kalkavan H, Green DR. MOMP, cell suicide as a BCL-2 family business. Cell Death & Differentiation 25, 46–55 (2018).

31. Dai H, Meng XW, Kaufmann SH. Mitochondrial apoptosis and BH3 mimetics. F1000Res 5, 2804 (2016).

32. Brumatti G, Kaloni D, Castro FA, Amarante-Mendes GP. BH3 mimetics and TKI combined therapy for Chronic Myeloid Leukemia. Biochemical Journal 480, 161–176 (2023).

33. Zhu Y, et al. Identification of a novel senolytic agent, navitoclax, targeting the Bcl-2 family of anti-apoptotic factors. Aging Cell 15, 428–435 (2016).

34. Tse C, et al. ABT-263: a potent and orally bioavailable Bcl-2 family inhibitor. Cancer Res 68, 3421–3428 (2008).

35. Godfraind JM, KrnjeviĆ K, Pumain R. Unexpected Features of the Action of Dinitrophenol on Cortical Neurones. Nature 228, 562–564 (1970).

36. Juthberg SK, Brismar T. Effect of metabolic inhibitors on membrane potential and ion conductance of rat astrocytes. Cell Mol Neurobiol 17, 367–377 (1997).

37. Buckler KJ, Vaughan-Jones RD. Effects of mitochondrial uncouplers on intracellular calcium, pH and membrane potential in rat carotid body type I cells. J Physiol 513 **(Pt** **3**), 819–833 (1998).

38. Triana-Martínez F, et al. Author Correction: Identification and characterization of Cardiac Glycosides as senolytic compounds. Nat Commun 11, 4771 (2020).

39. Zhu Y, et al. The Achilles’ heel of senescent cells: from transcriptome to senolytic drugs. Aging Cell 14, 644–658 (2015).

40. Zhu Y, et al. New agents that target senescent cells: the flavone, fisetin, and the BCL-X(L) inhibitors, A1331852 and A1155463. Aging (Albany NY) 9, 955–963 (2017).

41. Yousefzadeh MJ, et al. Fisetin is a senotherapeutic that extends health and lifespan. EBioMedicine 36, 18–28 (2018).

42. Stöckl P, et al. Partial uncoupling of oxidative phosphorylation induces premature senescence in human fibroblasts and yeast mother cells. Free Radic Biol Med 43, 947–958 (2007).

43. Kenwood BM, et al. Identification of a novel mitochondrial uncoupler that does not depolarize the plasma membrane. Molecular Metabolism 3, 114–123 (2014).

44. Axelrod CL, et al. BAM15-mediated mitochondrial uncoupling protects against obesity and improves glycemic control. EMBO Mol Med 12, e12088 (2020).

45. Alexopoulos SJ, et al. Mitochondrial uncoupler BAM15 reverses diet-induced obesity and insulin resistance in mice. Nature Communications 11, 2397 (2020).

46. Marroquin LD, Hynes J, Dykens JA, Jamieson JD, Will Y. Circumventing the Crabtree Effect: Replacing Media Glucose with Galactose Increases Susceptibility of HepG2 Cells to Mitochondrial Toxicants. Toxicological Sciences 97, 539–547 (2007).

47. Ewald JA, Desotelle JA, Wilding G, Jarrard DF. Therapy-induced senescence in cancer. J Natl Cancer Inst 102, 1536–1546 (2010).

48. Saleh T, et al. Therapy-Induced Senescence: An “Old” Friend Becomes the Enemy. Cancers (Basel) 12, 882 (2020).

49. Short S, Fielder E, Miwa S, von Zglinicki T. Senolytics and senostatics as adjuvant tumour therapy. EBioMedicine 41, 683–692 (2019).

50. Chen K, et al. Mitochondrial mutations and mitoepigenetics: Focus on regulation of oxidative stress-induced responses in breast cancers. Semin Cancer Biol 83, 556–569 (2022).

51. Bousset L, Gil J. Targeting senescence as an anticancer therapy. Mol Oncol 16, 3855–3880 (2022).

52. Kohli J, et al. Targeting anti-apoptotic pathways eliminates senescent melanocytes and leads to nevi regression. Nat Commun 13, 7923 (2022).

53. Troiani M, et al. Single-cell transcriptomics identifies Mcl-1 as a target for senolytic therapy in cancer. Nat Commun 13, 2177 (2022).

54. Aguayo-Mazzucato C, et al. Acceleration of β Cell Aging Determines Diabetes and Senolysis Improves Disease Outcomes. Cell Metab 30, 129–142.e124 (2019).

55. Bussian TJ, Aziz A, Meyer CF, Swenson BL, van Deursen JM, Baker DJ. Clearance of senescent glial cells prevents tau-dependent pathology and cognitive decline. Nature 562, 578–582 (2018).

56. Chang J, et al. Clearance of senescent cells by ABT263 rejuvenates aged hematopoietic stem cells in mice. Nat Med 22, 78–83 (2016).

57. Fielder E, et al. Short senolytic or senostatic interventions rescue progression of radiation-induced frailty and premature ageing in mice. eLife 11, e75492 (2022).

58. Lérida-Viso A, et al. Pharmacological senolysis reduces doxorubicin-induced cardiotoxicity and improves cardiac function in mice. Pharmacol Res 183, 106356 (2022).

59. Mylonas KJ, et al. Cellular senescence inhibits renal regeneration after injury in mice, with senolytic treatment promoting repair. Sci Transl Med 13, eabb0203 (2021).

60. Pan J, et al. Inhibition of Bcl-2/xl With ABT-263 Selectively Kills Senescent Type II Pneumocytes and Reverses Persistent Pulmonary Fibrosis Induced by Ionizing Radiation in Mice. Int J Radiat Oncol Biol Phys 99, 353–361 (2017).

61. Peng X, et al. Cellular senescence contributes to radiation-induced hyposalivation by affecting the stem/progenitor cell niche. Cell Death Dis 11, 854 (2020).

62. Walaszczyk A, et al. Pharmacological clearance of senescent cells improves survival and recovery in aged mice following acute myocardial infarction. Aging Cell 18, e12945 (2019).

63. Watanabe Y, et al. Navitoclax improves acute-on-chronic liver failure by eliminating senescent cells in mice. Hepatol Res 53, 460–472 (2023).

64. Fielder E, et al. Sublethal whole-body irradiation causes progressive premature frailty in mice. Mech Ageing Dev 180, 63–69 (2019).

65. Ross MF, et al. Lipophilic triphenylphosphonium cations as tools in mitochondrial bioenergetics and free radical biology. Biochemistry (Mosc) 70, 222–230 (2005).

66. Smith RA, Hartley RC, Murphy MP. Mitochondria-targeted small molecule therapeutics and probes. Antioxid Redox Signal 15, 3021–3038 (2011).

67. Smith RA, Hartley RC, Cochemé HM, Murphy MP. Mitochondrial pharmacology. Trends Pharmacol Sci 33, 341–352 (2012).

68. Chowdhury AR, Zielonka J, Kalyanaraman B, Hartley RC, Murphy MP, Avadhani NG. Mitochondria-targeted paraquat and metformin mediate ROS production to induce multiple pathways of retrograde signaling: A dose-dependent phenomenon. Redox Biology 36, 101606 (2020).

69. Hubackova S, et al. Selective elimination of senescent cells by mitochondrial targeting is regulated by ANT2. Cell Death & Differentiation 26, 276–290 (2019).

70. Kalyanaraman B, et al. Mitochondria-targeted metformins: anti-tumour and redox signalling mechanisms. Interface Focus 7, 20160109 (2017).

71. Saretzki G, Murphy MP, von Zglinicki T. MitoQ counteracts telomere shortening and elongates lifespan of fibroblasts under mild oxidative stress. Aging Cell 2, 141–143 (2003).

72. Dhanasekaran A, et al. Supplementation of endothelial cells with mitochondria-targeted antioxidants inhibit peroxide-induced mitochondrial iron uptake, oxidative damage, and apoptosis. J Biol Chem 279, 37575–37587 (2004).

73. Ly JD, Grubb DR, Lawen A. The mitochondrial membrane potential (deltapsi(m)) in apoptosis; an update. Apoptosis 8, 115–128 (2003).

74. Daouphars M, et al. Uncoupling of oxidative phosphorylation and Smac/DIABLO release are not sufficient to account for induction of apoptosis by sulindac sulfide in human colorectal cancer cells. Int J Oncol 26, 1069–1077 (2005).

75. Nair AB, Jacob S. A simple practice guide for dose conversion between animals and human. J Basic Clin Pharm 7, 27–31 (2016).

76. Childress ES, Alexopoulos SJ, Hoehn KL, Santos WL. Small Molecule Mitochondrial Uncouplers and Their Therapeutic Potential. Journal of Medicinal Chemistry 61, 4641–4655 (2018).

77. Dantas WS, et al. Mitochondrial uncoupling attenuates sarcopenic obesity by enhancing skeletal muscle mitophagy and quality control. J Cachexia Sarcopenia Muscle 13, 1821–1836 (2022).

78. Shrestha R, Johnson E, Byrne FL. Exploring the therapeutic potential of mitochondrial uncouplers in cancer. Mol Metab 51, 101222 (2021).

79. Farr JN, et al. Local senolysis in aged mice only partially replicates the benefits of systemic senolysis. J Clin Invest 133, e162519 (2023).

80. Budamagunta V, et al. Effect of peripheral cellular senescence on brain aging and cognitive decline. Aging Cell 22, e13817 (2023).

81. Petronilli V, Miotto G, Canton M, Colonna R, Bernardi P, Di Lisa F. Imaging the mitochondrial permeability transition pore in intact cells. Biofactors 8, 263–272 (1998).

82. Dimri GP, et al. A biomarker that identifies senescent human cells in culture and in aging skin in vivo. Proc Natl Acad Sci U S A 92, 9363–9367 (1995).

83. Di Veroli GY, et al. Combenefit: an interactive platform for the analysis and visualization of drug combinations. Bioinformatics 32, 2866–2868 (2016).

84. Cameron KM, Miwa S, Walker C, von Zglinicki T. Male mice retain a metabolic memory of improved glucose tolerance induced during adult onset, short-term dietary restriction. Longev Healthspan 1, 3 (2012).

85. Whitehead JC, et al. A clinical frailty index in aging mice: comparisons with frailty index data in humans. J Gerontol A Biol Sci Med Sci 69, 621–632 (2014).

86. Fielder E, et al. Anti-inflammatory treatment rescues memory deficits during aging in nfkb1−/− mice. Aging Cell 19, e13188 (2020).

