## Supplemental Figures and Table for "Mild uncoupling of mitochondria synergistically enhances senolytic specificity and sensitivity of BH3 mimetics"

**Supplementary Material**

**Supplementary Figures:**

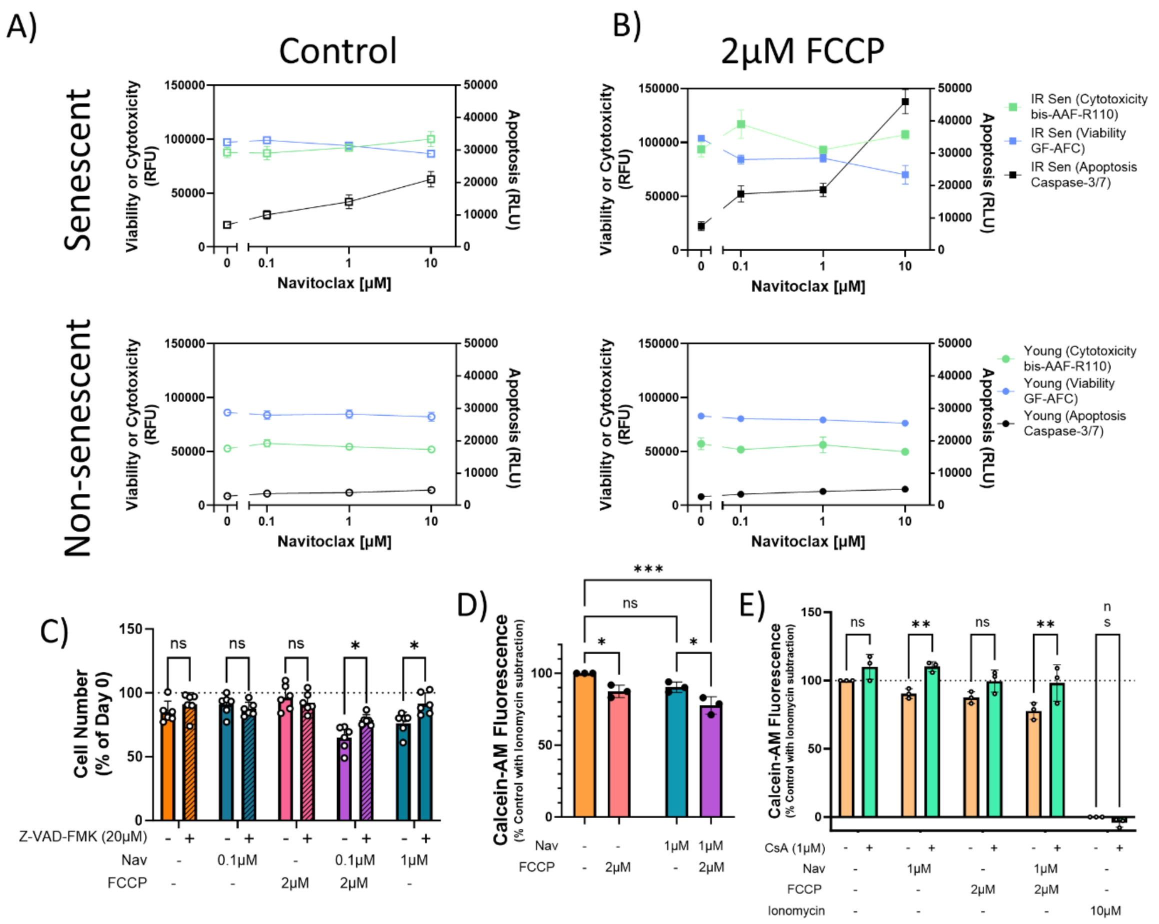


**Suppl. Fig S2.1: The combination of Navitoclax and FCCP enhances apoptosis of senescent HDFs.** A) and B) Cytotoxity (green), Viability (blue) and apoptosis induction (black) as measured in senescent (top) and non-senescent (bottom) fibroblasts under the indicated concentrations of Navitoclax without (A) and with (B) addition of 2µM FCCP. C) Senescent HDFs were treated with the indicated drugs +/- addition of the pan-caspase inhibitor z-vad-fmk for 24 hours, N=6. 2-way ANOVA with Šídák's multiple comparison test for the difference between control and z-VAD-FMK treatment of each condition, * p≤0.05. D) mPTP opening assay in senescent HDFs treated with indicated combination of drugs for 3 hours. Calcein-AM fluorescence is quenched by $CoCl_2$, which is normally impermeable to mitochondria. When the mPTP opens $CoCl_2$ can enter and quench Calcein-AM, leading to a decrease in signal. E) Action of the mPTP inhibitor CsA is then shown on each combination. Data are normalised to maximal release (by 10µM Ionomycin) and are from 3 independent experiments. 2-way ANOVA with Šídák's multiple comparison test, * p≤0.05, ** p≤0.005, *** p≤0.001

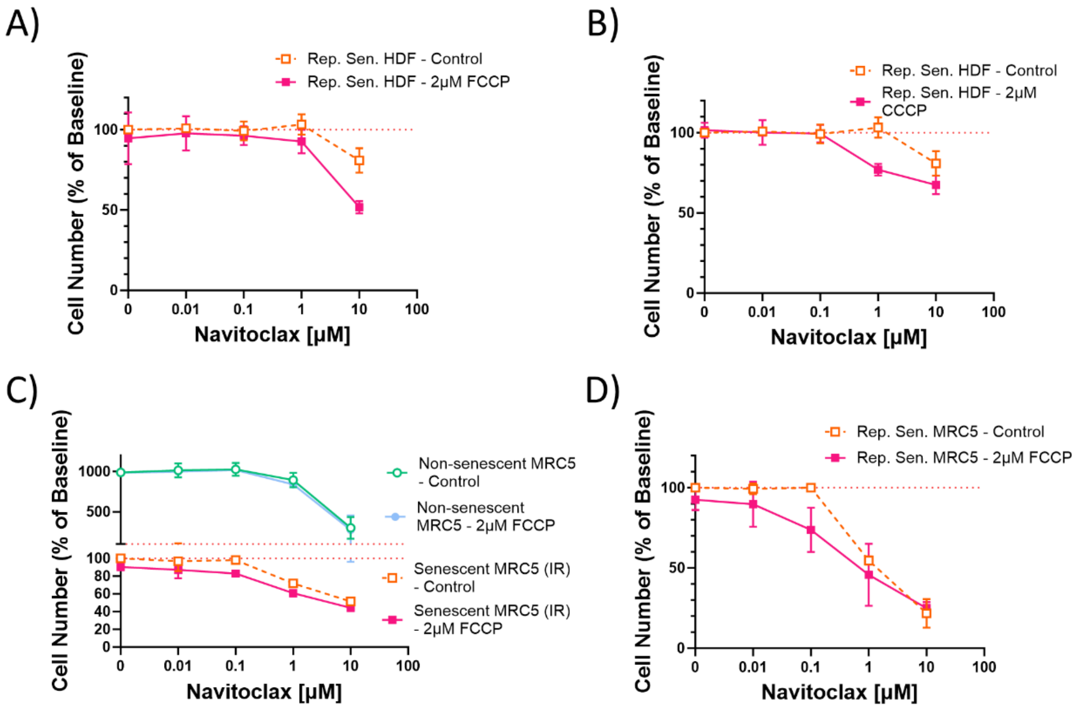


**Suppl. Fig. S2.2: Navitoclax/uncoupler combinations are effective in multiple models of senescence.** A-B) HDFs were grown to replicative senescence and treated with Navitoclax, with and without FCCP (A) and CCCP (B), respectively. C) Senescence was induced in MRC5 fibroblasts by 20Gy irradiation 10 days prior to treatment, or D) MRC5 fibroblasts were grown to replicative senescence and treated with Navitoclax, with and without FCCP. Dose response curves after 72 hours exposure to the drugs in normal media, N=3 independent experiments.

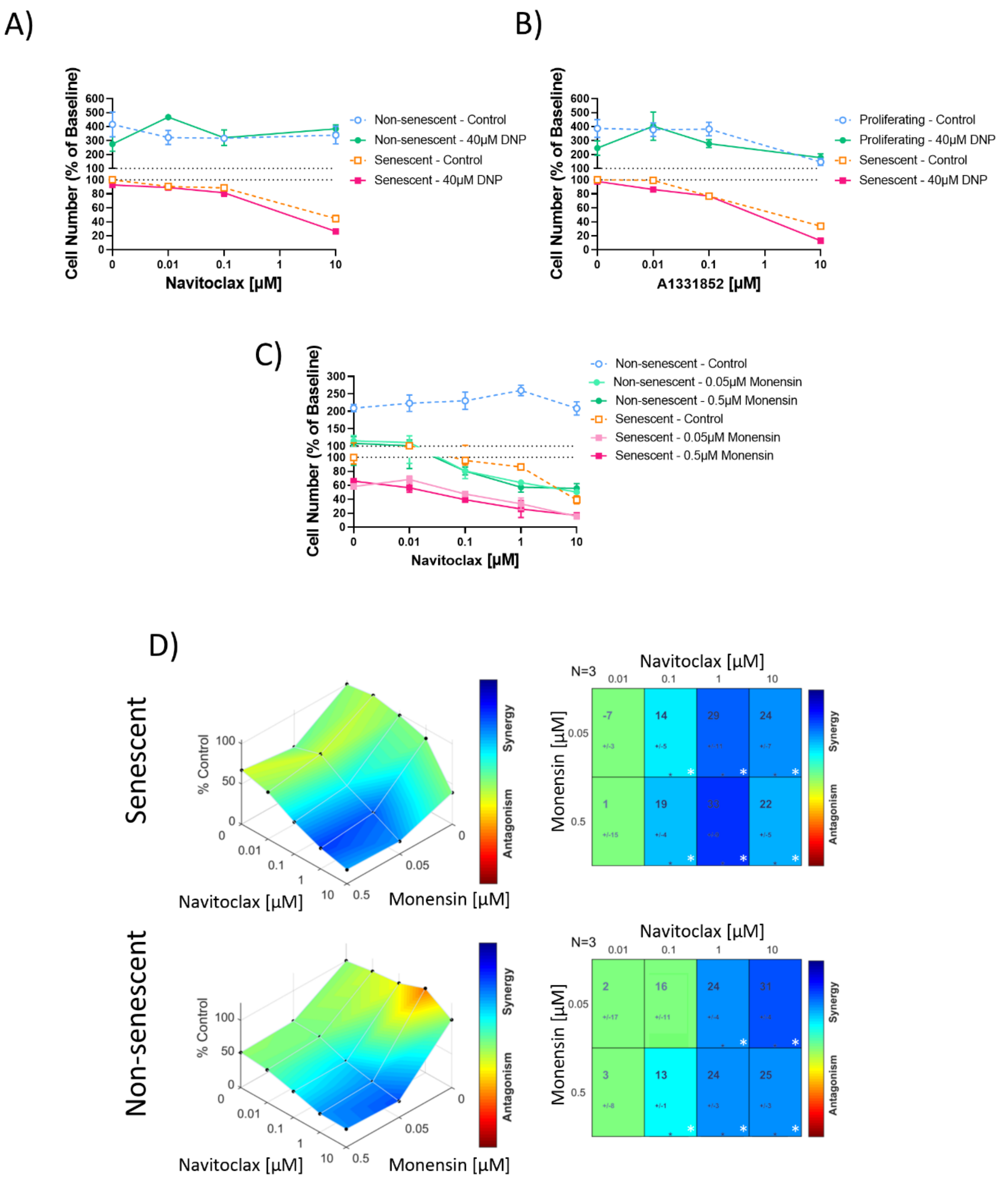


**Suppl. Fig. S2.3: No synergistic senolytic effects for combination of BH3 mimetics with a non-specific uncoupler or Monensin.** A)-C) Titration curves for HDFs with the indicated BH3 mimetic and either 40µM DNP (A) and B) or Monensin (C) in the indicated concentrations. Data are mean ± SEM, N=3. D) Highest Single Agent analysis showing cell viability compared to control with Navitoclax and Monensin in senescent (top) and non-senescent HDFs (bottom). Synergy/antagonism is shown by colour and, synergy score with change from expected value. * p≤0.05.

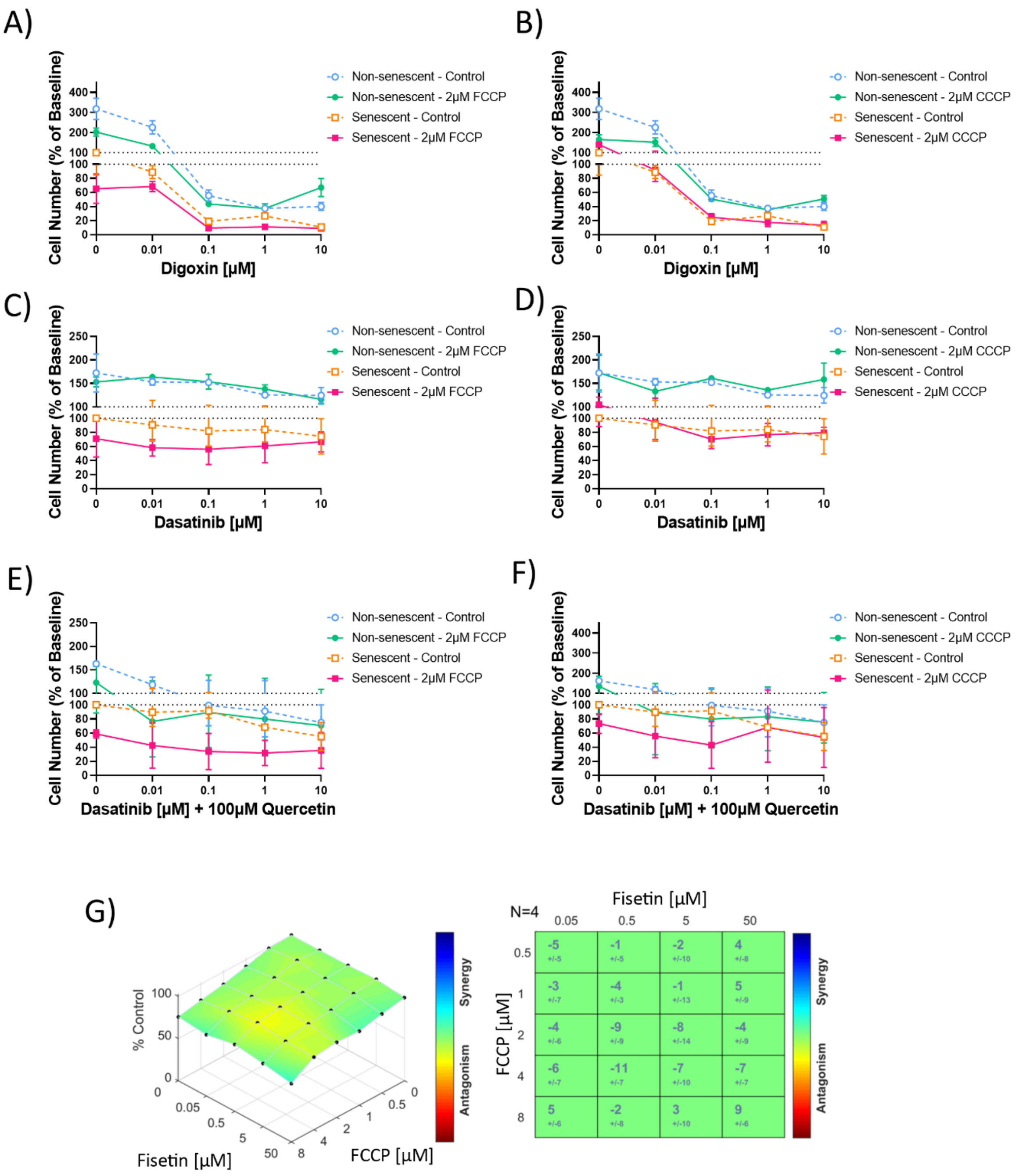


**Suppl. Fig. S2.4: Senolytics other than BH3 mimetics do not show synergism with mitochondrial uncouplers.** HDFs (non-senescent and in X-Ray induced senescence) were treated with the indicated combinations. A-B) Digoxin titration with either 2µM FCCP (A) or 2µM CCCP (B). C-D) Dasatinib titration with either 2µM FCCP (C) or 2µM CCCP (D). E-F) Titration with Dasatinib plus 100µM Quercetin with either 2µM FCCP (E) or 2µM CCCP (F). Dose response curves after 72 hours exposure to the drugs in normal media. Data are from 2 experimental repeats for Digoxin with mean ± SD shown, and mean ± SD from N=2 Independent experiments for Dasatinib plus Quercetin. G) Highest Single Agent analysis showing cell viability compared to control with Fisetin and FCCP in senescent HDFs with no significant change.

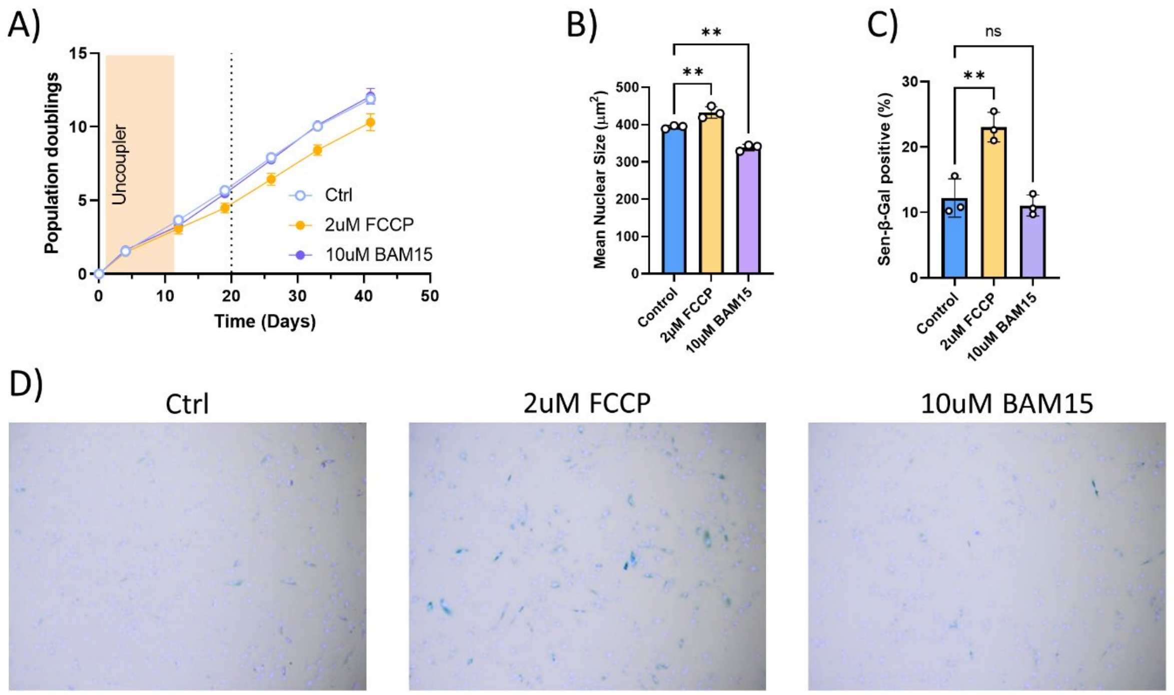


**Suppl. Fig S3.1: Long-term exposure to BAM15 does not induce senescence in HDFs.** A) Growth curves of HDFs continuously treated with the indicated uncouplers. B) Mean nuclear size of HDFs treated for 10 days with the indicated uncouplers. C) Fractions of Senescence-associated β-Galactosidase (Sen-β-Gal) positive HDFs treated for 10 days with the indicated uncouplers. Data are mean ± SEM, N=3. D) Representative images of Sen-β-Gal activity in HDFs treated with the indicated uncouplers.

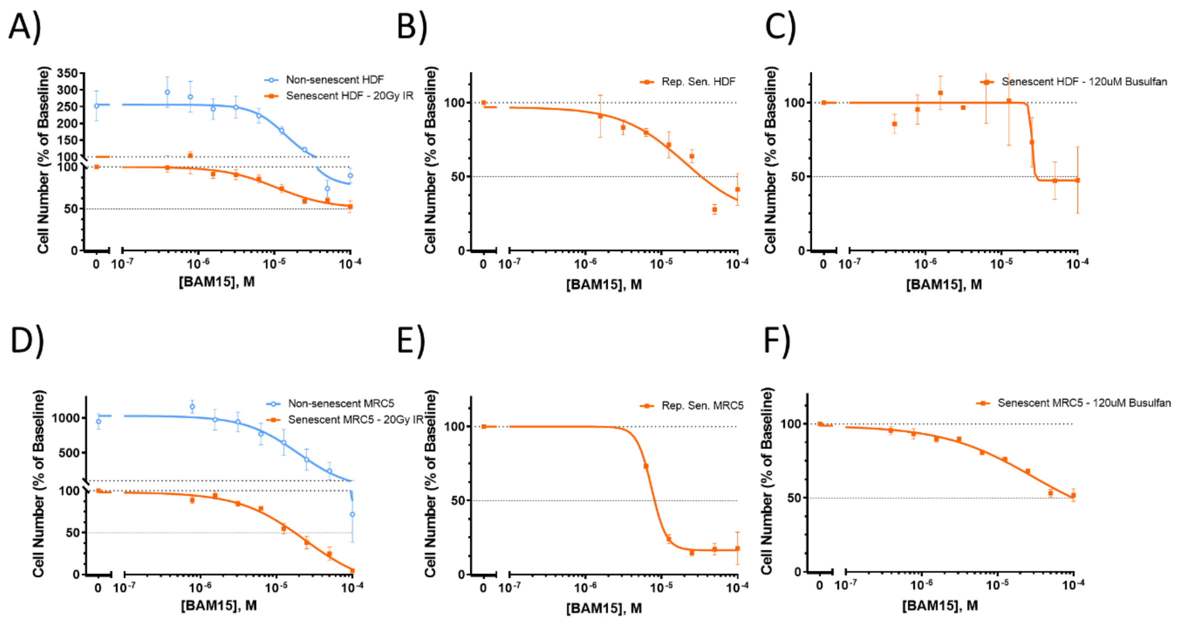

**Suppl. Fig S3.2: Effects of BAM15 alone on human fibroblasts.** Titration curves for HDFs (A) – C)) and MRC5 fibroblasts (D) – F)). Cells in X-Ray induced senescence and non-senescent cells (A) and D). Cells in replicative senescence (B) and E)). Cells in Busulfan-induced senescence (C) and F)). Data are mean ± SEM, N=2-6.

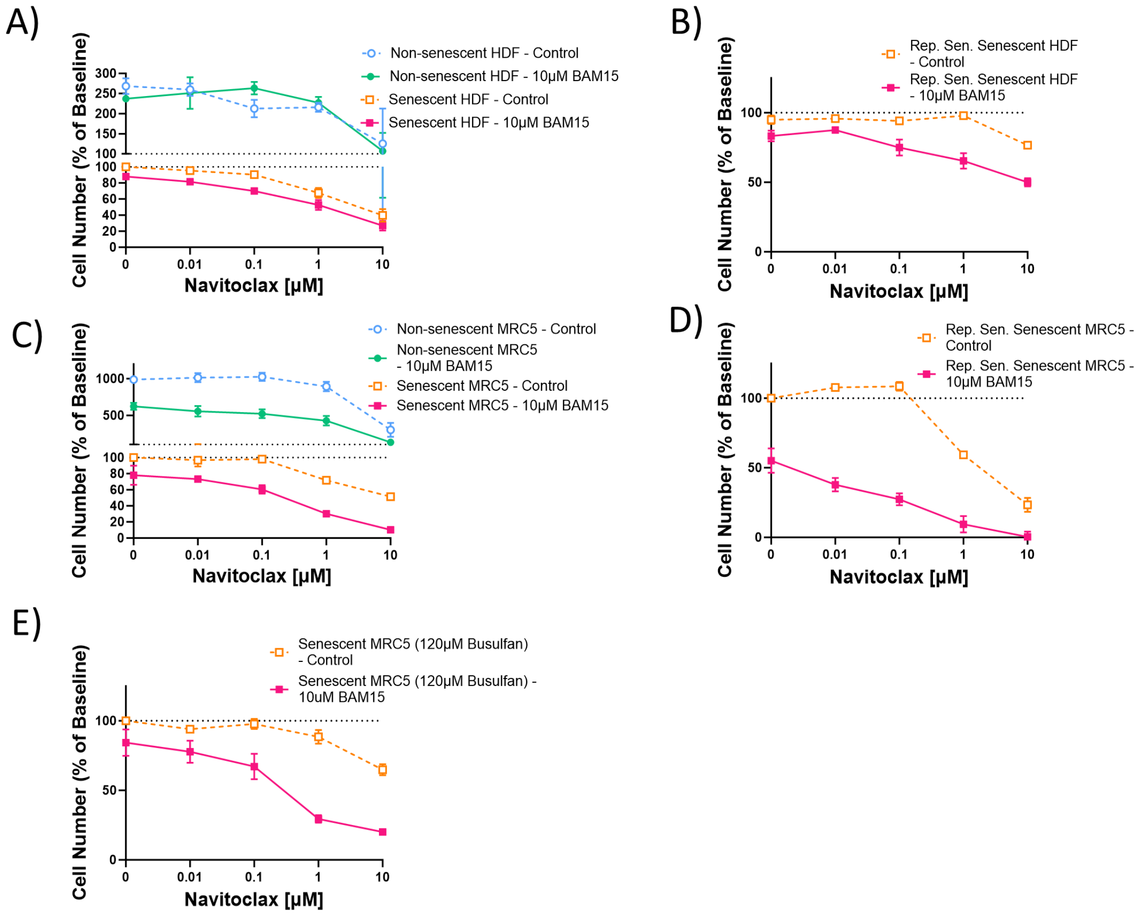


**Suppl. Fig S3.3: Navitoclax/BAM15 combinations are effective in multiple models of senescence.** Titration curves for Navitoclax treatment of A) stress-induced and B) replicatively senescent HDFs with and without BAM15. C-E) Titration curves for Navitoclax treatment of irradiation-induced (C), replicatively senescent (D) and chemotherapy induced (E) senescent MRC5 fibroblasts, with and without BAM15. Data are mean ± SEM from 2-7 experiments.

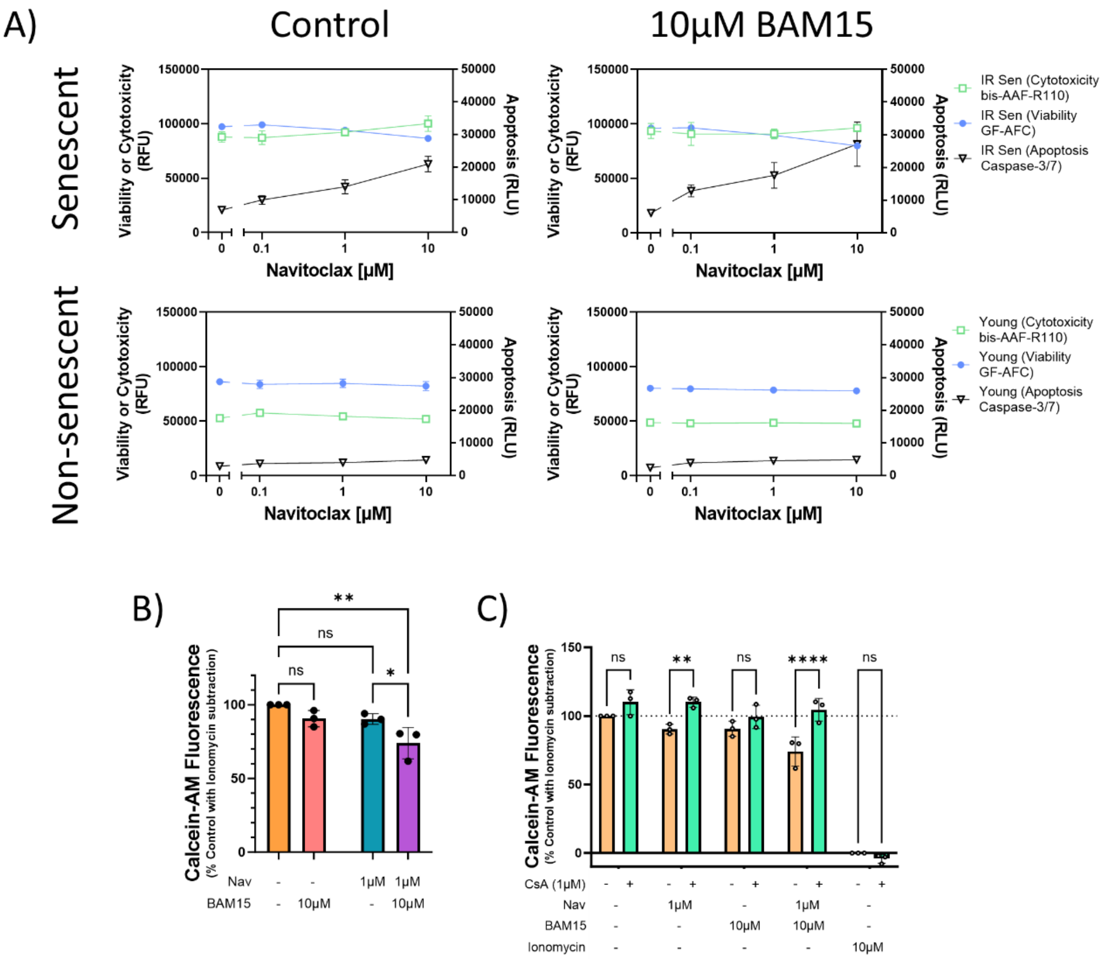


**Suppl. Fig S3.4: The combination of Navitoclax and BAM15 enhances apoptosis and induces mPTP pore opening in senescent HDFs.** A) Cytotoxity (green), Viability (blue) and apoptosis induction (black) as measured in senescent (top) and non-senescent (bottom) HDFs under the indicated concentrations of Navitoclax without (left) and with (right) addition of 10µM BAM15. Data are mean ± SEM, N=3. B) mPTP opening assay in senescent HDFs treated with the indicated combinations of drugs for 3 hours. C) Action of the mPTP inhibitor CsA is then shown on each combination in B), maximum release by 10µM Ionomycin is also shown which is not rescued by CsA (unlike 1µM Ionomycin, data not shown). Data are normalised to maximal release (by 10µM Ionomycin). 2-way ANOVA with Šídák's multiple comparison test, * p≤0.05, ** p≤0.005, **** p≤0.0001.

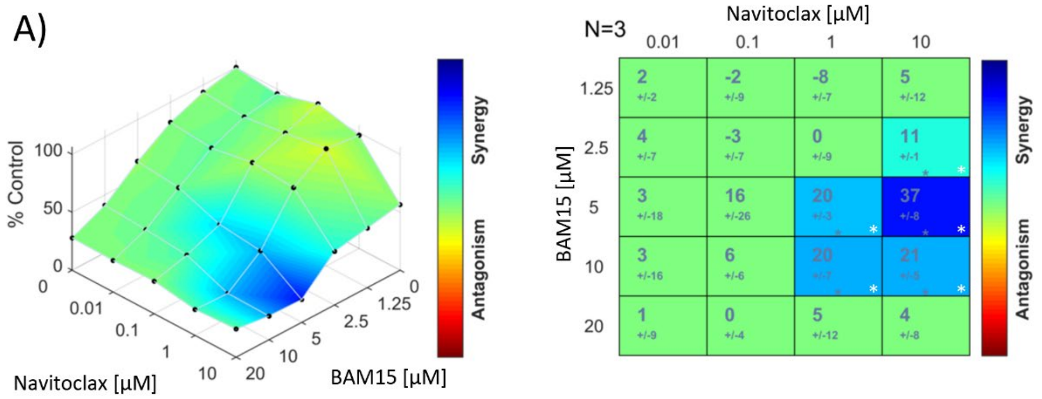


**Suppl. Fig S3.5: Forcing cellular bioenergetics to mitochondrial ATP generation by using Galactose media enhances the effect of Navitoclax and BAM15 combination.** A) Highest Single Agent analysis showing cell viability in senescent HDFs compared to control for Navitoclax and BAM15 in glucose free, galactose (5.5mM) containing media. Synergy/antagonism is shown by colour and, synergy score with change from expected value. N=3, *$p<0.05$. Peak synergistic concentration requires a lower dose of BAM15 compared to normal media (see Fig. 3A for comparison).

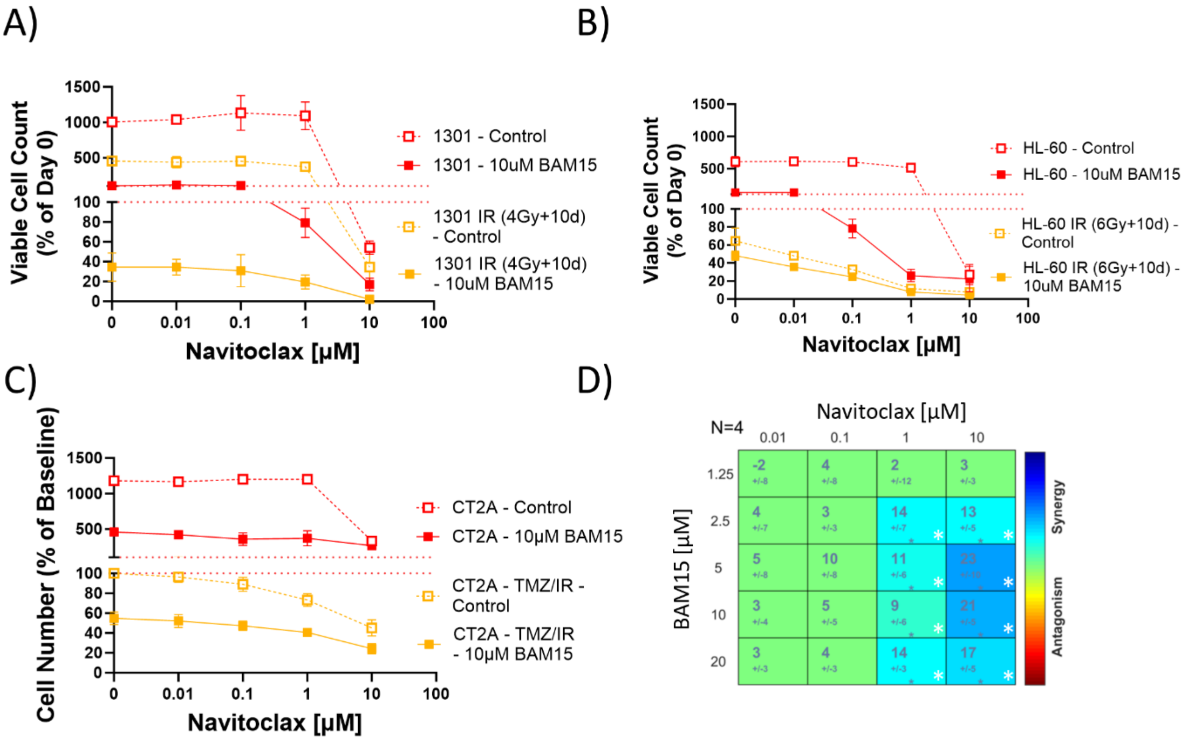


**Suppl. Fig S3.6: Combination with the uncoupler BAM15 synergistically enhances the killing efficacy of Navitoclax for both senescent and non-senescent tumour cells.** A) -C) Titration curves for non-senescent (red) and senescent (yellow) tumour cells (A: 1301; B: HL60, C: CT-2A) with (full symbols) and without (open symbols) 10µM BAM15. Senescence was induced by X-Ray irradiation (A, B) or irradiation and TMZ treatment (C). D) Highest Single Agent analysis showing cell viability compared to control with combinations of Navitoclax and BAM15 in senescent CT-2As. Synergy/antagonism is shown by colour and synergy score with change from expected value. N=4, *p<0.05.

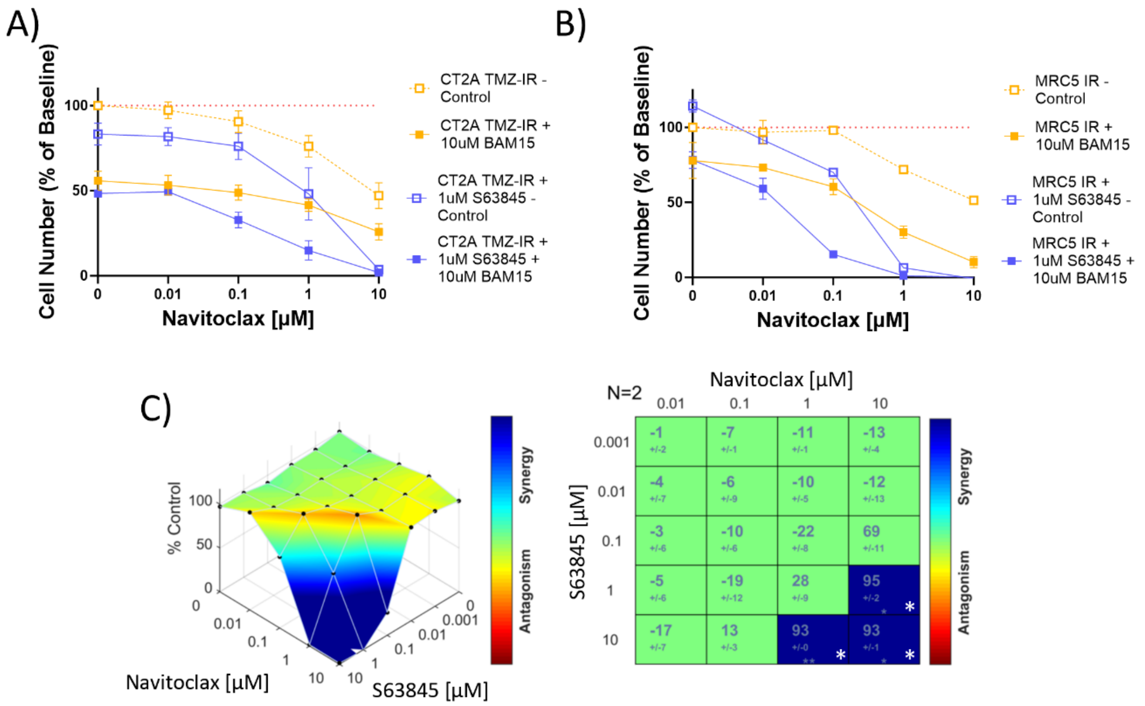


**Suppl. Fig S3.7: Mcl-1 inhibition further enhances the killing effect of the Navitoclax/BAM15 combination.** A-B) Titration curves for senescent CT-2A (A, senescence induction by irradiation plus TMZ) and MRC5 (B, senescence induction by irradiation) cells with (closed symbols) or without (open symbols) 10µM BAM15 and with (blue) or without (yellow) 1µM of the Mcl-1 inhibitor S63845. C) Highest Single Agent analysis showing cell viability compared to control with Navitoclax and S63845 in non-senescent HDFs. Synergy/antagonism is shown by colour and, synergy score with change from expected value. N=2, *p<0.05. Data are Mean ± SEM from 2-9 experiments.

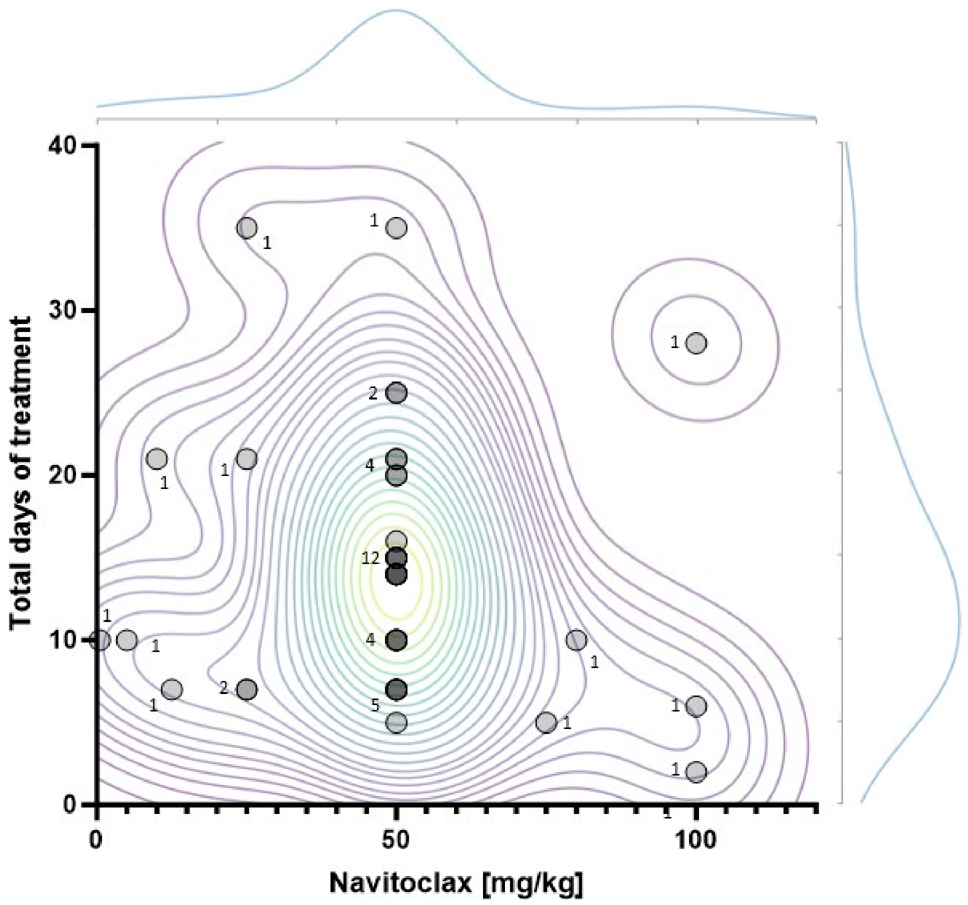


**Suppl. Fig S4.1:. Kernel Density Estimation plot of Navitoclax oral gavage dosing regimens in published studies.** A PubMed search was performed as ((Navitoclax) OR (ABT-263)) AND (mouse) and papers ascertaining to senescence, senolytics and ageing in mice were selected from this list for inclusion (last updated 2023-04-04). Total days of treatment shown on the Y-axis (distribution curve of results on top), with dose of Navitoclax by oral gavage on the X-axis (distribution curve of results on right) for each dosing regimen identified (Regimens and references are given in supplemental table 1). Number of separate experimental sets per data point on the graph are shown.

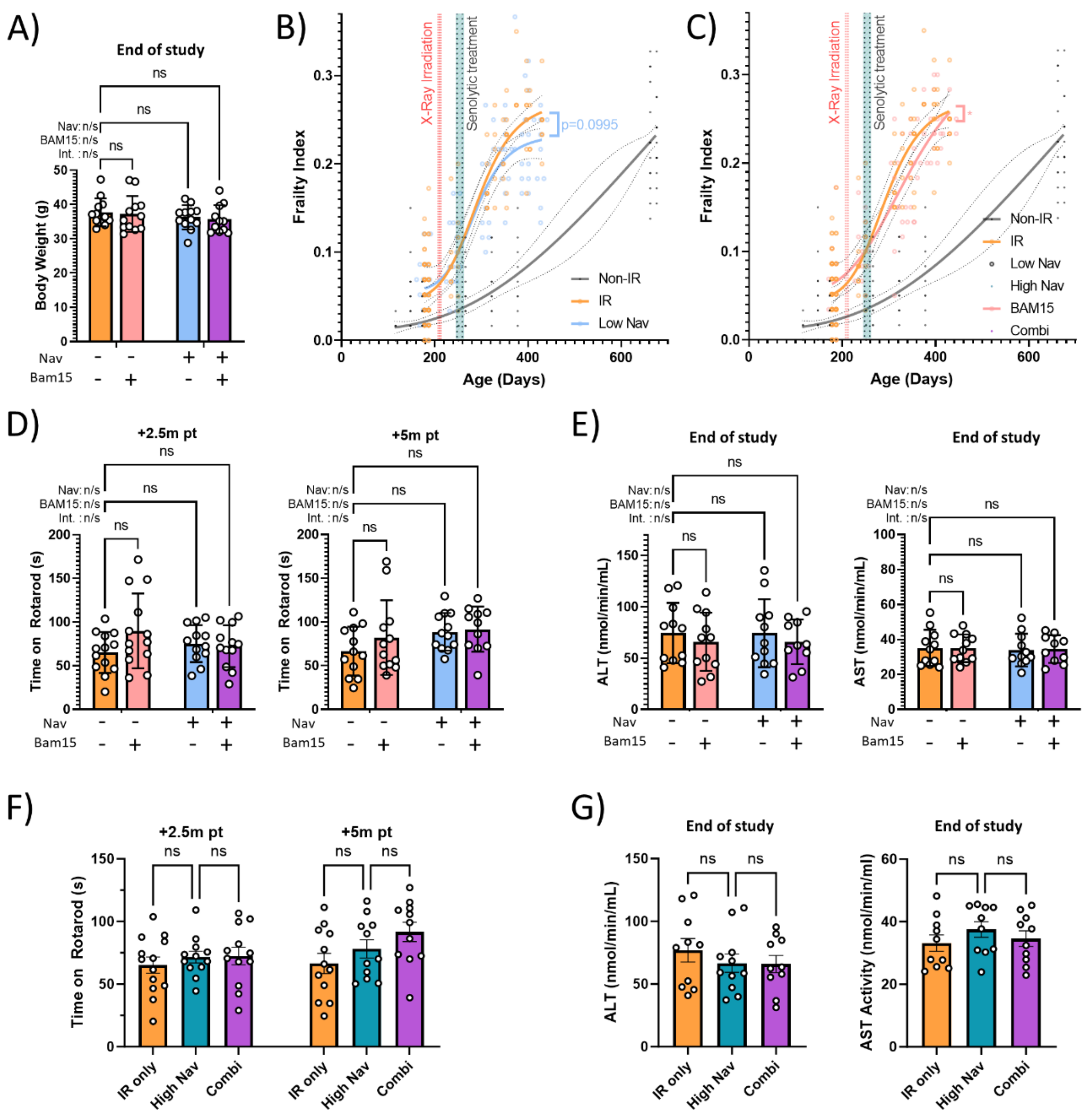


**Suppl. Fig S4.2: Further data on physiological effects of the combined treatment *in vivo*.** A) Body mass of irradiated mice treated with the indicated interventions at end of study. B) Impact of low Navitoclax alone (blue) on frailty. C) Impact of BAM15 alone (red) on frailty. Other data as in Figs. 4B and C. D) and F) Time to fall on a Rotarod for the indicated treatments at 2.5 months (left) and 5 months (right) past treatment. E) and G) Liver enzyme ALT (left) and AST (right) concentrations in serum of irradiated mice treated with the indicated interventions at end of study. Comparisons by 1- or 2-way ANOVA as appropriate with Šídák's multiple comparisons post-hoc test against the untreated irradiated control. N≥11.

**Supplementary Table S1: Published dosing regimens of Navitoclax as senolytic in mice**

| Dose/Day | Days | Cycles | Total Days | Total Dose | Paper | Reference |
|---|---|---|---|---|---|---|
| **100 mg/kg** | 28 | | 28 | 2800 mg/kg | Inhibition of antiapoptotic BCL-2 proteins with ABT-263 induces fibroblast apoptosis, reversing persistent pulmonary fibrosis | 1 |
| **100 mg/kg** | 6 | | 6 | 600 mg/kg | Senescence, Necrosis, and Apoptosis Govern Circulating Cell-free DNA Release Kinetics | 2 |
| **100 mg/kg** | 2 | | 2 | 200 mg/kg | Galacto-conjugation of Navitoclax as an efficient strategy to increase senolytic specificity and reduce platelet toxicity | 3 |
| **80 mg/kg** | 5 | 2x | 10 | 800 mg/kg | Senolytic-Mediated Elimination of Head and Neck Tumor Cells Induced Into Senescence by Cisplatin | 4 |
| **75 mg/kg** | 5 | | 5 | 375 mg/kg | Obesity triggers tumoral senescence and renders poorly immunogenic malignancies amenable to senolysis | 5 |
| **50 mg/kg** | 21 | | 21 | 1050 mg/kg | Eliminating Senescent Cells Can Promote Pulmonary Hypertension Development and Progression | 6 |
| **50 mg/kg** | 15 | | 15 | 750 mg/kg | Eliminating Senescent Cells Can Promote Pulmonary Hypertension Development and Progression | 6 |
| **50 mg/kg** | 5 | | 5 | 250 mg/kg | Navitoclax improves acute-on-chronic liver failure by eliminating senescent cells in mice | 7 |
| **50 mg/kg** | 7 | 2x | 14 | 700 mg/kg | Single-cell analysis of senescent epithelia reveals targetable mechanisms promoting fibrosis | 8 |
| **50 mg/kg** | 7 | | 7 | 350 mg/kg | Muscle injury induces a transient senescence-like state that is required for myofiber growth during muscle regeneration | 9 |
| **50 mg/kg** | 15 | | 15 | 750 mg/kg | Muscle injury induces a transient senescence-like state that is required for myofiber growth during muscle regeneration | 9 |
| **50 mg/kg** | 7 | 2x | 14 | 700 mg/kg | Renal inflamm-aging provokes intra-graft inflammation following experimental kidney transplantation | 10 |
| **50 mg/kg** | 2 | 8x | 16 | 800 mg/kg | Pharmacological senolysis reduces doxorubicin-induced cardiotoxicity and improves cardiac function in mice | 11 |
| **50 mg/kg** | 5 | 5x | 25 | 1250 mg/kg | A motor neuron disease mouse model reveals a non-canonical profile of senescence biomarkers | 12 |
| **50 mg/kg** | 5 | 4x | 20 | 1000 mg/kg | Extracellular Nicotinamide Phosphoribosyltransferase Is a Component of the Senescence-Associated Secretory Phenotype | 13 |

| | | | | | | |
|---|---|---|---|---|---|---|
| **50 mg/kg** | 5 | 2x | 10 | 500 mg/kg | Extracellular Nicotinamide Phosphoribosyltransferase Is a Component of the Senescence-Associated Secretory Phenotype | 13 |
| **50 mg/kg** | 5 | 3x | 15 | 750 mg/kg | Efficacy and limitations of senolysis in atherosclerosis | 14 |
| **50 mg/kg** | 25 | | 25 | 1250 mg/kg | Elimination of Radiation-Induced Senescence in the Brain Tumor Microenvironment Attenuates Glioblastoma Recurrence | 15 |
| **50 mg/kg** | 5 | 2x | 10 | 500 mg/kg | Brahma-related gene-1 promotes tubular senescence and renal fibrosis through Wnt/β-catenin/autophagy axis | 16 |
| **50 mg/kg** | 10 | | 10 | 500 mg/kg | Targeting senescent cells improves functional recovery after spinal cord injury | 17 |
| **50 mg/kg** | 7 | 2x | 14 | 700 mg/kg | Cellular senescence inhibits renal regeneration after injury in mice, with senolytic treatment promoting repair | 18 |
| **50 mg/kg** | 7 | 3x | 21 | 1050 mg/kg | Cellular senescence inhibits renal regeneration after injury in mice, with senolytic treatment promoting repair | 18 |
| **50 mg/kg** | 7 | | 7 | 350 mg/kg | ABT-263 enhanced bacterial phagocytosis of macrophages in aged mouse through Beclin-1-dependent autophagy | 19 |
| **50 mg/kg** | 7 | | 7 | 350 mg/kg | Senescence-associated secretory phenotype promotes chronic ocular graft-vs-host disease in mice and humans | 20 |
| **50 mg/kg** | 7 | | 7 | 350 mg/kg | BH3 mimetics selectively eliminate chemotherapy-induced senescent cells and improve response in TP53 wild-type breast cancer | 21 |
| **50 mg/kg** | 7 | 2x | 14 | 700 mg/kg | Cellular senescence contributes to radiation-induced hyposalivation by affecting the stem/progenitor cell niche | 22 |
| **50 mg/kg** | 5 | 3x | 15 | 750 mg/kg | FBP1 loss disrupts liver metabolism and promotes tumorigenesis through a hepatic stellate cell senescence secretome | 23 |
| **50 mg/kg** | 14 | | 14 | 700 mg/kg | The Senolytic Drug Navitoclax (ABT-263) Causes Trabecular Bone Loss and Impaired Osteoprogenitor Function in Aged Mice | 24 |
| **50 mg/kg** | 5 | 4x | 20 | 1000 mg/kg | Acceleration of β Cell Aging Determines Diabetes and Senolysis Improves Disease Outcomes | 25 |
| **50 mg/kg** | 7 | 2x | 14 | 700 mg/kg | Pharmacological clearance of senescent cells improves survival and recovery in aged mice following acute myocardial infarction | 26 |
| **50 mg/kg** | 5 | 7x | 35 | 1750 mg/kg | Clearance of senescent glial cells prevents tau-dependent pathology and cognitive decline | 27 |
| **50 mg/kg** | 5 | 2x | 10 | 500 mg/kg | Inhibition of Bcl-2/xl With ABT-263 Selectively Kills Senescent Type II Pneumocytes and Reverses Persistent Pulmonary Fibrosis Induced by Ionizing Radiation in Mice | 28 |
| **50 mg/kg** | 7 | 2x | 14 | 700 mg/kg | Clearance of senescent cells by ABT263 rejuvenates aged hematopoietic stem cells in mice | 29 |

| | | | | | | |
|---|---|---|---|---|---|---|
| **25 mg/kg** | 21 | | 21 | 525 mg/kg | Eliminating Senescent Cells Can Promote Pulmonary Hypertension Development and Progression | 6 |
| **25 mg/kg** | 5 | 7x | 35 | 875 mg/kg | Gestational arsenite exposure augments hepatic tumors of C3H mice by promoting senescence in F1 and F2 offspring via different pathways | 30 |
| **25 mg/kg** | 7 | | 7 | 175 mg/kg | Tissue damage and senescence provide critical signals for cellular reprogramming in vivo | 31 |
| **25 mg/kg** | 7 | | 7 | 175 mg/kg | Senescence-associated secretory phenotype promotes chronic ocular graft-vs-host disease in mice and humans | 20 |
| **12.5 mg/kg** | 7 | | 7 | 87.5 mg/kg | Senescence-associated secretory phenotype promotes chronic ocular graft-vs-host disease in mice and humans | 20 |
| **10 mg/kg** | 21 | | 21 | 210 mg/kg | Eliminating Senescent Cells Can Promote Pulmonary Hypertension Development and Progression | 6 |
| **5 mg/kg** | 5 | 2x | 10 | 50 mg/kg | Short senolytic or senostatic interventions rescue progression of radiation-induced frailty and premature ageing in mice | 32 |

Supplementary references:


1. Cooley JC, et al. Inhibition of antiapoptotic BCL-2 proteins with ABT-263 induces fibroblast apoptosis, reversing persistent pulmonary fibrosis. JCI Insight 8, e163762 (2023).

2. Rostami A, Lambie M, Yu CW, Stambolic V, Waldron JN, Bratman SV. Senescence, Necrosis, and Apoptosis Govern Circulating Cell-free DNA Release Kinetics. Cell Rep 31, 107830 (2020).

3. González-Gualda E, et al. Galacto-conjugation of Navitoclax as an efficient strategy to increase senolytic specificity and reduce platelet toxicity. Aging Cell 19, e13142 (2020).

4. Ahmadinejad F, et al. Senolytic-Mediated Elimination of Head and Neck Tumor Cells Induced Into Senescence by Cisplatin. Mol Pharmacol 101, 168-180 (2022).

5. Fournier F, et al. Obesity triggers tumoral senescence and renders poorly immunogenic malignancies amenable to senolysis. Proc Natl Acad Sci U S A 120, e2209973120 (2023).

6. Born E, et al. Eliminating Senescent Cells Can Promote Pulmonary Hypertension Development and Progression. Circulation 147, 650-666 (2023).

7. Watanabe Y, et al. Navitoclax improves acute-on-chronic liver failure by eliminating senescent cells in mice. Hepatol Res 53, 460-472 (2023).

8. O'Sullivan ED, et al. Single-cell analysis of senescent epithelia reveals targetable mechanisms promoting fibrosis. JCI Insight 7, e154124 (2022).

9. Young LV, et al. Muscle injury induces a transient senescence-like state that is required for myofiber growth during muscle regeneration. Faseb j 36, e22587 (2022).

10. He A, et al. Renal inflamm-aging provokes intra-graft inflammation following experimental kidney transplantation. Am J Transplant 22, 2529-2547 (2022).

11. Lérida-Viso A, et al. Pharmacological senolysis reduces doxorubicin-induced cardiotoxicity and improves cardiac function in mice. Pharmacol Res 183, 106356 (2022).

12. Torres P, et al. A motor neuron disease mouse model reveals a non-canonical profile of senescence biomarkers. Dis Model Mech 15, dmm049059 (2022).

13. Kuehnemann C, et al. Extracellular Nicotinamide Phosphoribosyltransferase Is a Component of the Senescence-Associated Secretory Phenotype. Front Endocrinol (Lausanne) 13, 935106 (2022).

14. Garrido AM, et al. Efficacy and limitations of senolysis in atherosclerosis. Cardiovasc Res 118, 1713-1727 (2022).

15. Fletcher-Sananikone E, et al. Elimination of Radiation-Induced Senescence in the Brain Tumor Microenvironment Attenuates Glioblastoma Recurrence. Cancer Res 81, 5935-5947 (2021).

16. Gong W, et al. Brahma-related gene-1 promotes tubular senescence and renal fibrosis through Wnt/β-catenin/autophagy axis. Clin Sci (Lond) 135, 1873-1895 (2021).

17. Paramos-de-Carvalho D, et al. Targeting senescent cells improves functional recovery after spinal cord injury. Cell Rep 36, 109334 (2021).

18. Mylonas KJ, et al. Cellular senescence inhibits renal regeneration after injury in mice, with senolytic treatment promoting repair. Sci Transl Med 13, eabb0203 (2021).

19. Zhang Y, Tang LH, Lu J, Xu LM, Cheng BL, Xiong JY. ABT-263 enhanced bacterial phagocytosis of macrophages in aged mouse through Beclin-1-dependent autophagy. BMC Geriatr 21, 225 (2021).

20. Yamane M, et al. Senescence-associated secretory phenotype promotes chronic ocular graft-vs-host disease in mice and humans. Faseb j 34, 10778-10800 (2020).

21. Shahbandi A, et al. BH3 mimetics selectively eliminate chemotherapy-induced senescent cells and improve response in TP53 wild-type breast cancer. Cell Death Differ 27, 3097-3116 (2020).

22. Peng X, et al. Cellular senescence contributes to radiation-induced hyposalivation by affecting the stem/progenitor cell niche. Cell Death Dis 11, 854 (2020).

23. Li F, et al. FBP1 loss disrupts liver metabolism and promotes tumorigenesis through a hepatic stellate cell senescence secretome. Nat Cell Biol 22, 728-739 (2020).

24. Sharma AK, et al. The Senolytic Drug Navitoclax (ABT-263) Causes Trabecular Bone Loss and Impaired Osteoprogenitor Function in Aged Mice. Front Cell Dev Biol 8, 354 (2020).

25. Aguayo-Mazzucato C, et al. Acceleration of β Cell Aging Determines Diabetes and Senolysis Improves Disease Outcomes. Cell Metab 30, 129-142.e124 (2019).

26. Walaszczyk A, et al. Pharmacological clearance of senescent cells improves survival and recovery in aged mice following acute myocardial infarction. Aging Cell 18, e12945 (2019).

27. Bussian TJ, Aziz A, Meyer CF, Swenson BL, van Deursen JM, Baker DJ. Clearance of senescent glial cells prevents tau-dependent pathology and cognitive decline. Nature 562, 578-582 (2018).

28. Pan J, et al. Inhibition of Bcl-2/xl With ABT-263 Selectively Kills Senescent Type II Pneumocytes and Reverses Persistent Pulmonary Fibrosis Induced by Ionizing Radiation in Mice. Int J Radiat Oncol Biol Phys 99, 353-361 (2017).

29. Chang J, et al. Clearance of senescent cells by ABT263 rejuvenates aged hematopoietic stem cells in mice. Nat Med 22, 78-83 (2016).

30. Okamura K, Suzuki T, Nohara K. Gestational arsenite exposure augments hepatic tumors of C3H mice by promoting senescence in F1 and F2 offspring via different pathways. Toxicol Appl Pharmacol 408, 115259 (2020).

31. Mosteiro L, et al. Tissue damage and senescence provide critical signals for cellular reprogramming in vivo. Science 354, aaf4445 (2016).

32. Fielder E, et al. Short senolytic or senostatic interventions rescue progression of radiation-induced frailty and premature ageing in mice. eLife 11, e75492 (2022).
